# Visualizing Synaptic Dopamine Efflux with a 2D Nanofilm

**DOI:** 10.1101/2022.01.19.476937

**Authors:** Chandima Bulumulla, Andrew T. Krasley, Deepika Walpita, Abraham G. Beyene

## Abstract

Chemical neurotransmission constitutes one of the fundamental modalities of communication between neurons. Monitoring release of these chemicals has traditionally been difficult to carry out at spatial and temporal scales relevant to neuron function. To understand chemical neurotransmission more fully, we need to improve the spatial and temporal resolutions of measurements for neurotransmitter release. To address this, we engineered a chemi-sensitive, two-dimensional nanofilm that facilitates subcellular visualization of the release and diffusion of the neurochemical dopamine with synaptic resolution, quantal sensitivity, and simultaneously from hundreds of release sites. Using this technology, we were able to monitor the spatiotemporal dynamics of dopamine release in dendritic processes, a poorly understood phenomenon. We found that dopamine release is broadcast from a subset of dendritic processes as hotspots that have a mean spatial spread of ≈3.2 µm (full width at half maximum) and are observed with a mean spatial frequency of 1 hotspot per ≈7.5 µm of dendritic length. Major dendrites of dopamine neurons and fine dendritic processes, as well as dendritic arbors and dendrites with no apparent varicose morphology participated in dopamine release. Remarkably, these release hotspots colocalized with Bassoon, suggesting that Bassoon may contribute to organizing active zones in dendrites, similar to its role in axon terminals.

## Introduction

Chemical neurotransmission generally falls under one of two broad categories: fast synaptic transmission or neuromodulation. Synapses that mediate rapid communication between most excitatory and inhibitory synapses in the brain primarily employ glutamate or γ-aminobutyric acid (GABA). Such chemical communication occurs at highly specialized synaptic structures that have nanoscale spatial organization and operate with millisecond temporal precision.^1–3^ In contrast, neuromodulators, including biogenic amines, neuropeptides, and hormones, operate at different spatiotemporal scales. Neuromodulatory synapses do not exhibit a close apposition to their partners but act on receptors that are extrasynaptically localized, and signal through G-protein coupled intracellular mechanisms.^4^ This suggests that neuromodulators diffuse from their release sites in adequate quantities to influence target receptors.^5^

Generally, the nature of the chemical synapse is less well understood for neuromodulators, which differ from their classical counterparts not just in their mechanisms of action on receptors but also in their secretory apparatus and spatiotemporal dynamics. For dopamine, one of the most important neuromodulators in the brain, the challenge is compounded by certain unique features pertinent to dopamine neurobiology. Dopaminergic neurons are known for their large size, and exhibit highly ramified axonal arborizations and dense varicosities.^6–8^ Studies have sought to establish the molecular determinants of dopamine release from these dense axonal arbors, but these attempts do not rely on measurement of dopamine efflux with single release site resolution.^9, 10^ Previous studies have demonstrated that dopamine neurons possess the machinery for co-release of other neurotransmitters, including glutamate and GABA, but to what extent, if any, co-release events spatially overlap remains insufficiently understood.^11–15^ Electrochemical and microdialysis assays in midbrain regions have shown that somatodendritic release of dopamine constitutes an important component of dopamine signaling, with notable implications in disease and behavior.^16–21^ However, the spatial and temporal dynamics of dendritic release events and their regulatory mechanisms remain poorly characterized. In sum, there is a pressing need for tools with appropriate sensitivity, kinetics, and sub-cellular spatial resolution to explore mechanisms of neurochemical release, including that of dopamine.

In this work, we developed an assay that facilitates visualization of the efflux of dopamine from active zones with synaptic resolution and quantal sensitivity. Here, we define efflux to mean the two-dimensional (2D) broadening of released dopamine quanta. In axonal arbors, we observe that dopamine release arises from a sparse set of varicosities, and we are able to assign the observed spatially defined effluxes to individually identified boutons. In dendrites and dendritic arbors of dopamine neurons, we similarly visualized spatially resolved effluxes of dopamine from putative dendritic active zones, which have been less well understood than release zones in axons. Our results show that dopamine neurons can sustain robust levels of release from their dendritic processes. Indeed, we demonstrate that dendritic release is fast and Ca^2+^ dependent, suggesting gating of dopamine release by a Ca^2+^-mediated release machinery, reminiscent of classical active zones. Dopamine efflux was observed at major dendrites, fine dendritic processes and at junctions of the soma and dendrites but rarely directly from the cell-body itself. Soma of dopamine neurons were observed to receive strong dopaminergic input due to release and diffusion from proximal dendrites. Retrospective super resolution imaging at identified release sites shows that Bassoon, long established as a presynaptic scaffolding protein of the cytomatrix in presynaptic active zones, is also enriched at dopamine release sites in dendrites and dendritic arbors. The expression of vesicular SNARE protein synaptobrevin-2 correlated with dopamine release activity in dendritic segments, establishing its functional utility in dopamine neurons. Therefore, active zones in dendrites appear to utilize a constellation of presynaptic and SNARE-complex proteins that are responsible for coordinating release, similar to those observed in classical synapses. This study offers a technology that facilitates visualization of the spatial and temporal efflux of dopamine and deploys the technology to shed light on somatodendritic dopamine release, a facet of dopaminergic neurobiology that has heretofore been poorly characterized.

## Results

### Visualizing active zone dopamine efflux from axons and dendrites

To a first order approximation, once released from a synaptic active zone, the temporal evolution of the released chemical should approximate that of diffusion from a point source, characterized by an isotropic expansion from the point of release but constrained by transporter activity and local three-dimensional ultrastructure. Such a signal can only be fully measured if the sensing platform sufficiently samples the underlying signal in both the spatial and temporal domains. However, the inability of current technologies to measure chemical efflux sufficiently in the spatial domain limits our ability to study chemical synapses. We addressed this challenge by using near-infrared fluorescent dopamine nanosensors to image single dopamine release sites from rat primary mid-brain neurons. The nanosensors are assembled from oligonucleotide-functionalized, single wall carbon nanotubes in solution phase^22, 23^ and have previously been used to image dopamine release in striatal acute slices and cultured cells but were lacking in synaptic information.^24, 25^ In this study, we drop-cast glass coverslips with dopamine nanosensors to produce a 2D layer of a turn-on fluorescent, dopamine-sensitive surface that can effectively image dopamine diffusion from a point source (Figure 1A). Temporally, the sensors exhibit millisecond-scale turn-on responses, which enabled real-time imaging of dopamine’s temporal evolution. We named the engineered surface DopaFilm, a 2D engineered film that affords video-rate filming of dopamine spatiotemporal dynamics with subcellular spatial and millisecond temporal resolution.

**Figure 1:**
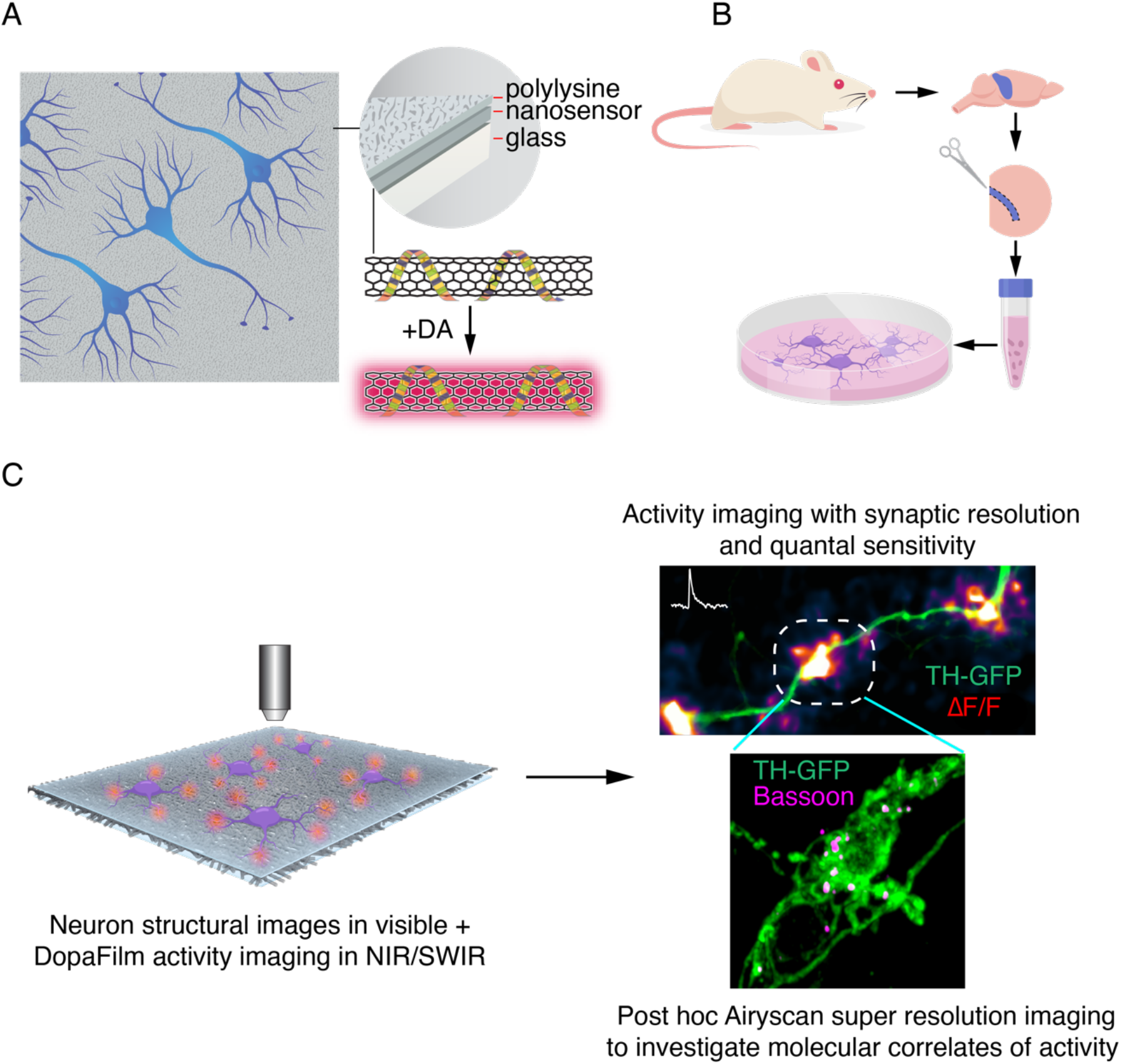
Schematic of DopaFilm imaging protocol. (A) Schematic of DopaFilm. (B) Workflow for preparing dopamine neuron primary cultures from the rat mid brain regions highlighted in blue. Neurons are grown on dishes with an engineered, chemi-sensitive, and fluorescent surface (DopaFilm) at the bottom. (C) Imaging DopaFilm fluorescence activity in cultured dopamine neurons. Immunocytochemistry and Airyscan super resolution imaging are carried out following DopaFilm activity imaging.

DopaFilm was validated by co-culturing rat primary mid-brain dopamine neurons with cortical neurons on the nanofilm surface for up to 6 weeks (typical period for an experiment) (Figures 1B, 1C). Dopamine neurons in culture exhibited stereotyped morphology, with thick major dendrites arising from the soma that go on to ramify into dendritic arbors, and axonal arbors that branch and ramify on a scale of millimeters from a single neuron (Figure S1). DopaFilm fluoresced in the near-infrared (NIR) to short-wave infrared (SWIR) regions of the spectrum (850-1350 nm) when imaged with a 785 nm excitation laser, permitting its multiplexed deployment with existing optical technologies with no spectral overlap (Figure S2). The surface exhibited an isotropic turn-on response when exposed to exogenous dopamine wash, suggesting uniform sensor coverage and response (Figure S2A). Dopamine wash experiments revealed DopaFilm has the sensitivity to detect 1 nM concentrations, remained stable over the duration of a typical experiment, and has an apparent dissociation constant (*K_d_*) of 268 nM (Figure S2B, S2C). This compares to EC_50_ values of ∼1 µM for D_1_-like and ∼10 nM for D_2_-like dopamine receptors, suggesting that DopaFilm is sensitive to chemical secretions that have physiological relevance.^26^ In order to leverage DopaFilm’s advantageous photophysical properties, we developed a custom microscope that is optimized for broad spectrum imaging in the visible, NIR and SWIR regions of the spectrum (400-1400 nm), with integrated widefield and laser scanning confocal capabilities. The optimizations in the NIR and SWIR regions facilitated imaging and recording of activity with exceedingly high signal to noise (SNR) ratios, attaining SNRs in the range of 5 – 50 for most experiments. In this study, most imaging experiments were carried out in widefield epifluorescence mode using a 40x/0.8 NA objective (N40X-NIR, Nikon). This gave us a field of view (FOV) of 180 µm by 230 µm. In axonal arbors, the FOV contained several hundred dopaminergic varicosities, whereas in cell-body regions, we could simultaneously image activity around the soma, major dendrites, and dendritic arbors. Post hoc immunofluorescence super resolution images were collected on Zeiss LSM 880 with Airyscan mode.

We asked if dopamine neurons grown on DopaFilm can be evoked to release dopamine, and whether DopaFilm fluorescence transients can recapitulate the predicted spatiotemporal evolution of dopamine efflux. To drive this effort, we virally co-expressed TH-GFP (GFP expressed under the control of the rat Tyrosine Hydroxylase (TH) promoter) and Syn-ChrimsonR-tdTomato (the red shifted opsin, ChrimsonR, expressed under the control of synapsin promoter and fused to tdTomato for visualization). This co-expression paradigm facilitated identification and optical stimulation of putative dopamine neurons. 79% of TH-GFP+ neurons were confirmed to be dopaminergic in retrospective immunofluorescence against tyrosine hydroxylase, the rate limiting enzyme in dopamine biosynthesis. On the other hand, 100% of TH-immunoreactive neurons expressed the TH-GFP transgene that was delivered virally. Optimized seeding density allowed us to record activities arising from isolated dopamine neurons where no other neurons in the vicinity of the neuron of interest were TH+. We used 561 nm LED (5 pulses, 25 Hz) to depolarize dopaminergic neurons. A subset of dopamine neurons exhibited spontaneous spiking activity and we applied no external stimuli in those cases. Furthermore, most dopamine neurons exhibited stochastic release events that appeared to be action potential independent, and those events were also included in our data. All imaging experiments were carried out 3-4 weeks post viral infections and neurons were 4-5-weeks *in vitro* at time of activity imaging.

We first imaged in axonal arbors, where dopamine release is relatively better characterized through microdialysis and voltammetry measurements.^10, 27–29^ Whenever targeted to TH+ neurons (further confirmed by retrospective immunofluorescence experiments), our stimulation protocol elicited robust fluorescence transients from DopaFilm (Figure 2A, 2B, Movie S1) and the fluorescence hotspots co-localized with TH+ boutons in axonal arbors (Figure 2C). We observed diffusive broadening of the fluorescence hotspots in subsequent imaging frames (Figure 2A, *+1s*), and fluorescence transients returned to baseline in the post stimulation epoch (Figure 2A *post*, Figure 2B). The observed spatiotemporal evolution of DopaFilm hotspots is consistent with that of release and diffusion from multiple point-like sources localized in a 2D plane, with estimated mass diffusivities of ≈ (1.1 ± 0.8) x 10^-6^ cm^2^ s^-1^ (mean ± SD), in reasonable agreement with estimated values of diffusion coefficient for dopamine (Figure 2A, 2C).^30^ Despite the high density of axonal varicosities in the FOV, fluorescence transients were observed to emanate from a subset of varicosities while another subset of varicosities produced no corresponding ΔF/F fluorescence hotspots (Figure 2C, red arrows). This sparse-release observation is in agreement with results from previous studies.^10, 31^ DopaFilm hotspot activities were also observed to co-localize with dendrites of dopamine neurons (Figure 2D, 2E) and were additionally noted to arise from dendritic processes that comingled with the soma (Figure 2F, 2G). Stimulation of ChrimsonR+ but TH-neurons did not elicit fluorescence transient in DopaFilm, and no evoked nor spontaneous activities were noted in neurons that lacked TH-immunoreactivity despite being TH-GFP+ in the viral expression paradigm (Figure S3). Both evoked and spontaneous DopaFilm fluorescence transients were absent when imaging in extracellular Ca^2+^-free media (Figure S4).

**Figure. 2:**
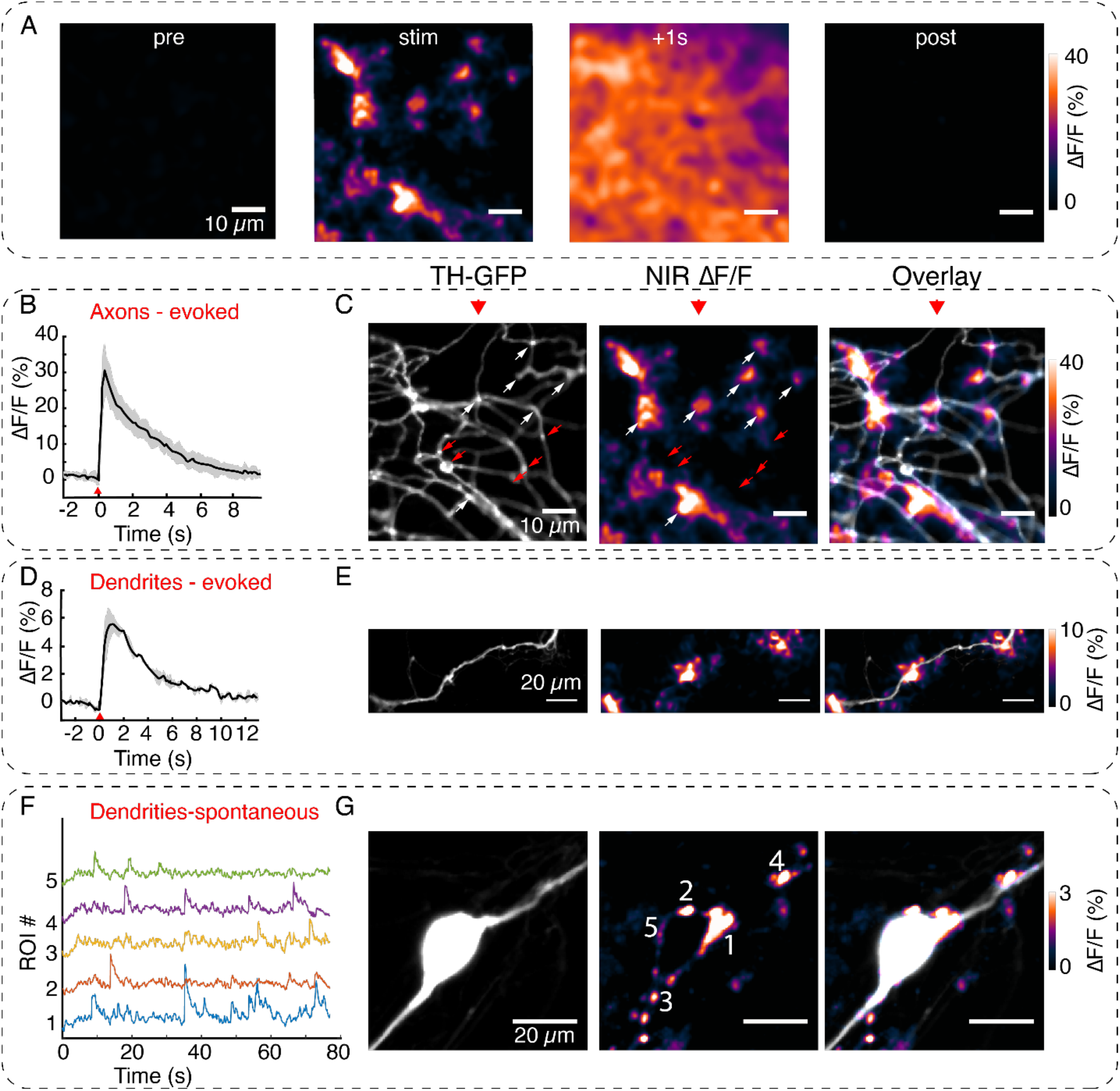
Visualizing active zone dopamine efflux from axons and dendrites. (A) ΔF/F images before stimulation (pre), at time of stimulation (stim), 1 second after stimulation (+ 1s) and after return to baseline (post). (B) Mean ± SD of ΔF/F traces from the imaging FOV in (A) averaged over n = 5 repeat stimulation runs. (C) TH-GFP image of axonal arbor and NIR ΔF/F image shown in (A) and overlay. The NIR ΔF/F frame corresponds to ‘stim’ and before diffusive broadening of the hotspots. (D, E) TH-GFP and ΔF/F activity from a dendrite of a dopamine neuron and overlay. Cell body not shown. Activity traces from dendrite are averaged over n = 3 stimulations. (F, G) Spontaneous activity from dendrites around cell body of a dopamine neuron and maximum intensity projection of the ΔF/F stack and overlay. (F) shows ΔF/F activity traces from regions of interest (ROI) numbered in (G). Red wedges in (B) and (D)= time of optical stimuli.

### DopaFilm hotspots localize to defined varicosities

We next explored the consistency of DopaFilm ΔF/F hotspot dynamics over multiple stimulation epochs and asked if repeat stimulations in the same FOV generated DopaFilm hotspots that localize to the same set of boutons. We carried out imaging in a FOV where we applied the same optical stimuli to drive hotspot activity over separate imaging sessions. In axon terminals, we observed that repeat stimulations can be carried out in a FOV up to n = 10 times (possibly longer) with little apparent depletion in activity, with rest periods of ∼2-3 minutes between stimuli (Figure 3A, 3B). We then computed the intensity-weighted centroid of each DopaFilm hotspot and compared the centroids across multiple stimulation repeats. We found that hotspot centroids were remarkably consistent across repeat stimulations (Figure 3C). To determine if the hotspots localized to the same set of varicosities, we compared DopaFilm hotspot centroids with the centroid of a TH+ varicosity that spatially overlapped with the DopaFilm hotspot from the overlay image (Figure 3B). We defined dopamine varicosities as puncta where TH-GFP intensity is at least 3x the mean intensity of TH-GFP expression along the process and computed the centroids of these TH+ boutons (Figure 3B center, green puncta, Figure 3D). Using an arbitrarily chosen origin as a reference point (0,0), we compared centroids of DopaFilm hotspots and TH+ boutons. Our results showed that these centroids matched with remarkable consistency across stimuli (no offset in some ROIs, <5 camera pixels for all ROIs, equivalent to <1.7 µm) (Figure 3E). These experiments demonstrate that DopaFilm activity hotspots can be faithfully localized to the same set of boutons across stimulation epochs and are therefore likely driven by the efflux of dopamine from putative active zones of these boutons. The ability to localize synaptic dopamine efflux to specific boutons is unique to this study and to our knowledge has not been demonstrated before.

**Figure 3:**
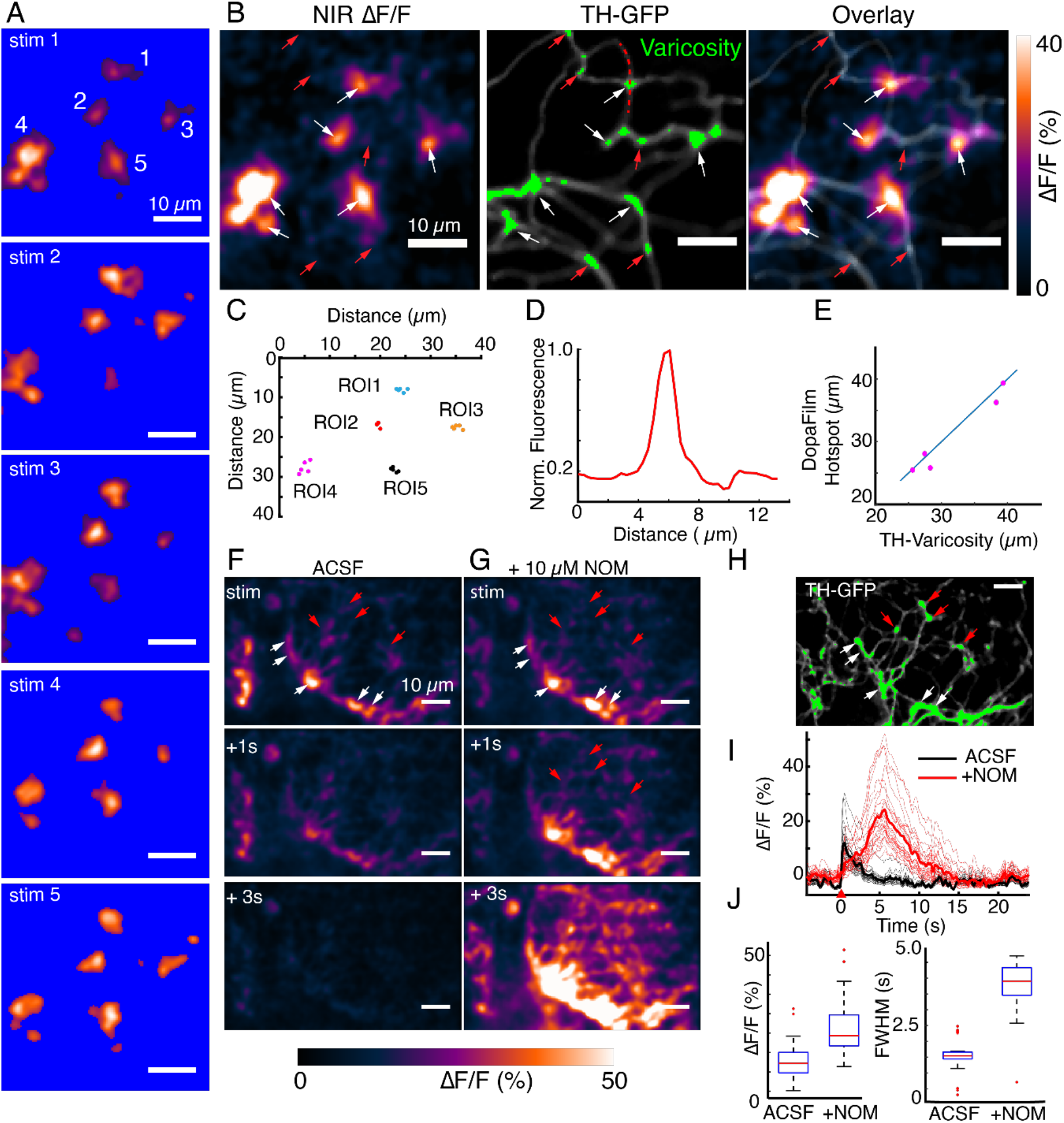
DopaFilm hotspots localize to defined varicosities. (A) Repeat optical stimulation produces a consistent set of DopaFilm hotspots (n=5 of n=10 stimulations shown), thresholded for better visualization. (B) Hotspot colocalization with TH-GFP varicosities (green) for stim #1. Green colors indicate GFP mean intensity > 3x the mean intensity along the process. (C) Centroid of hotpots (ROI1 to ROI5). Each data point of an ROI corresponds to one evoked imaging run and there are five total per ROI. (D) Line profile for one of the varicosities shown in (B). Line profile is calculated along the red curve shown in (B). (E) Parity plot for TH and DopaFilm hotspot centroids. Centroid of TH varicosities and averaged centroid of DopaFilm hotspots are plotted. Distance is calculated from the top left corner (=origin) of NIR ΔF/F image shown in B. (F, G) Spatiotemporal dynamics of imaging in ACSF and ACSF+nomifensine, respectively. (H) TH-GFP image of FOV in (F, G). Red arrows = no release. Scale bar = 10 µm (I) ΔF/F traces of hotspots in ACSF (black) and ACSF + 10 µM nomifensine (red) and their mean traces in bold. (J) Box plots comparing the effect of nomifensine on peak ΔF/F (left) and FWHM (right) of traces shown in (I). Unpaired t-test, *p* < 10^-4^ for both ΔF/F and FWHM data. See Methods for box plot definitions.

It is notable that not all TH-GFP+ boutons produced a corresponding DopaFilm activity (Figure 2C, Figure 3B, red arrows indicate no activity) despite appearing to satisfy the morphological criteria for a varicosity. We considered the possibility that the failure to detect activity from some putatively silent boutons was the result of rapid dopamine clearance, before DopaFilm detection. To test if dopamine clearance was critical to ‘silent’ boutons, we applied saturating levels of nomifensine (> 10 µM), a dopamine-specific reuptake inhibitor. Application of nomifensine dramatically slowed down the clearance of dopamine at previously active varicosities (mean full width at half maximum (FWHM) of 1.85 s before and 4.45 s after application of nomifensine) and increased the peak amplitude of the ΔF/F traces (mean peak ΔF/F = 15.7% before and 24.5% after application of nomifensine) (Figures 3F, 3G, 3I, 3J). Importantly, application of nomifensine did not reveal any subthreshold activity at silent varicosities that could have gone undetected in pre-nomifensine imaging sessions (Figure 3F, 3G). Thus, we conclude that DopaFilm hotspot activity arises from dopamine release at varicosities, and that the absence of DopaFilm fluorescence transient is likely an indication of a lack of dopamine release at release-incompetent varicosities. When coupled with the fact that DopaFilm activity is not observed in TH-cells, and that activity is absent when imaging in Ca^2+^-free media, we conclude that DopaFilm ΔF/F activity is a consequence of release of endogenous dopamine from active release sites. Nomifensine manipulation of the temporal dynamics of dopamine release provides additional evidence that DopaFilm possesses the kinetics necessary to recapitulate the dynamic behavior of dopamine release, diffusion and clearance.

### DopaFilm detects quantal release of dopamine

We next sought to establish the limit of detection of DopaFilm. In *in vitro* experiments, we determined that DopaFilm exhibits high sensitivity to dopamine and can detect 1 nM concentrations in bath application experiments (Figure S2). This suggested that DopaFilm may be sensitive enough to detect single events of quantized dopamine efflux from release sites. To determine the limit of detection from a practical sense, we carried out imaging experiments in a field of an axonal arbor of a dopamine neuron before and after bath application of tetrodotoxin (TTX), which inhibits action potential driven, synchronous neurotransmitter release while sparing stochastic and spontaneous release events. We first imaged activity in artificial cerebrospinal fluid (ACSF), our normal imaging buffer and then applied TTX (Figure 4A). As expected, bath application of 10 µM of TTX abolished evoked release of dopamine but stochastic fluorescence transients persisted (Figure 4A, 4B, 4C, Movie S2 before TTX, Movie S3 after TTX). The spatial extent of DopaFilm fluorescence hotspots were smaller after application of TTX (Figure 4A, center-vs-right panels, 4B, 4D, Figure S5), the peak amplitude of transients were diminished, and transients returned to baseline faster (Figure 4C, 4E, Figure S5). Notably, DopaFilm fluorescence transients in TTX were still detected with robust SNR of more than 5, where SNR is defined as the ratio of *ΔF* to the standard deviation of *F_0_* (Figure 4C, Figure S5). Incidentally, SNR for most of our TTX-free, optical stimulus-driven imaging in axons frequently exceeded 30. In dense axonal arbors where DopaFilm fluorescence hotspots could not be assigned to specific varicosities, application of TTX allowed visualization of spatial evolution from putative single release sites by eliminating release from neighboring active zones (Figure 4A, Figure S5). A frequency histogram of the peak amplitudes of DopaFilm transients in TTX fits a set of Gaussian curves whose spacings are approximately integer multiples, suggesting that DopaFilm fluorescence transients in TTX are driven by a quantized biological process (Figure 4F). In sum, our results demonstrate that DopaFilm has the sensitivity to detect quantal release events of dopamine and, to our knowledge, offers the only experimental observation of putative single-vesicle fusion events in a manner that is resolved in both spatial and temporal domains.

**Figure 4:**
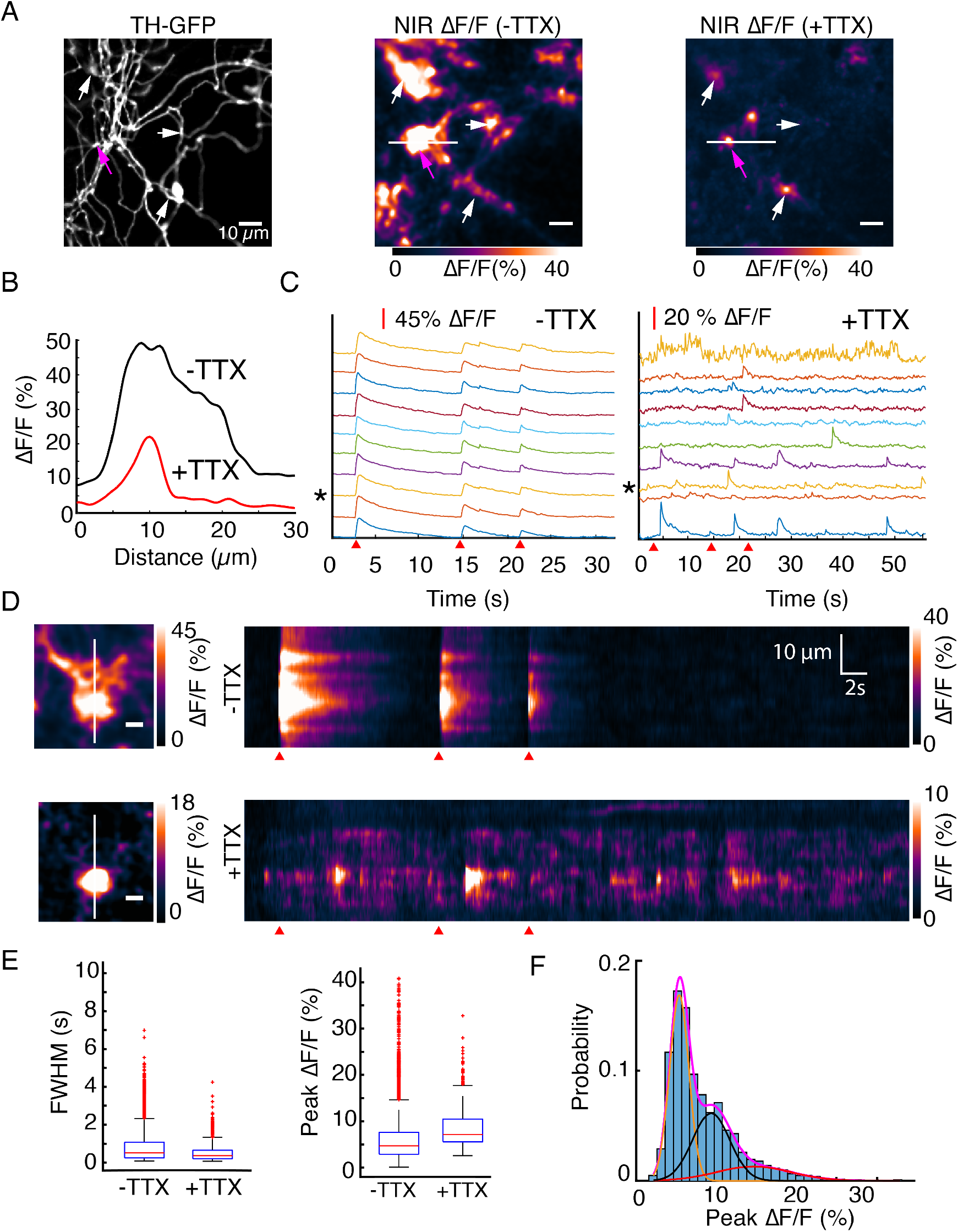
DopaFilm detects quantal release of dopamine. (A) TH-GFP image from an axonal arbor of a dopamine neuron (soma not shown). Peak ΔF/F of DopaFilm fluorescence transients with no TTX in imaging media (-TTX) and max ΔF/F projection of stack with 10 µM TTX in media (+ TTX). Arrows provided to aid comparison across FOVs. (B) ΔF/F profiles across the white lines depicted in (A). (C) Fluorescence transient traces for hotspots with (-TTX) and (+TTX), respectively. * indicates trace corresponding to magenta arrow in (A). Red wedges in (C) and (D) indicate times of optical stimuli. (D) Kymographs along the white lines shown in (A). Scale bar = 5 µm. (E) Box plots of FWHM (s) and ΔF/F amplitude (%) for (-TTX) and (+TTX) cases. Note that evoked and spontaneous data are pooled for -TTX. Unpaired t-test, *p* < 10^-4^ for both ΔF/F and FWHM data. See Methods for box plot definitions. (F) Frequency histogram of ΔF/F amplitudes for TTX data shown in (C) and Gaussian fits to the experimental data. Magenta = sum of three Gaussian curve fits. Means: µ_1_ = 5.7%, µ_2_ = 9.6%, µ_3_ = 15.1%.

### DopaFilm detects dopamine release activity from dendritic processes

To further demonstrate the utility of DopaFilm, we deployed it to study a poorly understood phenomena in dopamine neurobiology. Previous studies have demonstrated that dopamine is released from somatodendritic processes of dopamine neurons in the midbrain.^16–21^ Despite decades of research however, the release of dopamine at somata and dendrites of dopamine neurons remains incompletely understood.^32^ Because microdialysis and electrochemical assays, which have historically been employed to measure dopamine, do not have subcellular spatial resolution, it is still not clear if dopamine release detected in perikarya of dopamine neurons emanates from soma or dendrites. This has necessitated the use of the term ‘somatodendritic release’.^21^ Absence of small synaptic vesicles and synaptic morphologies in ultrastructural studies of dopamine neuron dendritic arbors has made the functional role of dendritic projections in mid-brain regions less obvious.^33–36^ We reasoned that DopaFilm could unravel some of these mysteries owing to its subcellular spatial and millisecond temporal resolution, and quantal sensitivity. When deployed for imaging activity in perikarya of dopamine neurons, DopaFilm detected both evoked and spontaneous transients of dopamine that clearly emanated from MAP2+ and TH+ dendritic processes of dopamine neurons (Figure 5, Figure S6). DopaFilm transients were observed at dendritic varicosities (i.e., bouton-like structures on dendrites) (Figure 5B) and dendritic processes that did not appear to have any varicose morphology (Figure 6A, 6G, Figure S7, Figure S8). From a spatial perspective, release at dendrites often resembled the localized, hot spot like activities seen when imaging in axonal arbors (Figure 5, Figure 6, Figure S7, Figure S10, Figure S11, Movie S4 for evoked, Movie S5 for spontaneous). In some dendritic processes, we observed activities that emanate from clustered active zones in contiguous segments of dendrites, producing hotspots that spread along the entire profile of the dendritic process (Figure 6G, Figure S8, Figure S10A bottom row, Movie S5). To rule out the possibility that what we perceived to be dendritic release could arise from commingling axonal processes in the same dendritic arbor, in all our images, we amplified the TH-GFP signal with an anti-GFP antibody and performed post hoc Airyscan super resolution imaging. We observed no axons that tracked with the dendrites and DopaFilm transients clearly localized to MAP2-positive and TH-GFP-positive processes, confirming their dendritic nature (Figure 5C, 5E, Figure 6H, Figure S6, Figure S8). DopaFilm ΔF/F peak amplitudes were smaller in dendrites compared to those measured from axonal release sites and transients returned to baseline faster (Figure S9A, S9B). We observed a notable difference in the spatial propagation propensity of dopamine release in axonal arbors vs. dendrites. In dendritic processes, DopaFilm hotspot activity remained confined to the vicinity of release site, which contrasted with the more diffusive dynamics observed in axons (Figure S9C, S9D, S9E). We computed the FWHM of each hotspot as a proxy to estimate the spatial spread of dopamine from a dendritic release site and obtained values of ≈3.2 µm ± 3 µm (mean ± SD). In axons, we observed spatial spreads of ≈6.6 µm ± 3.6 µm (mean ± SD) (Figure S9E). Dendritic hotspots were encountered at a frequency of ≈7.5 µm ± 0.7 µm (mean ± SD) along the active dendrites which participated in release (Figure S10).

**Figure 5:**
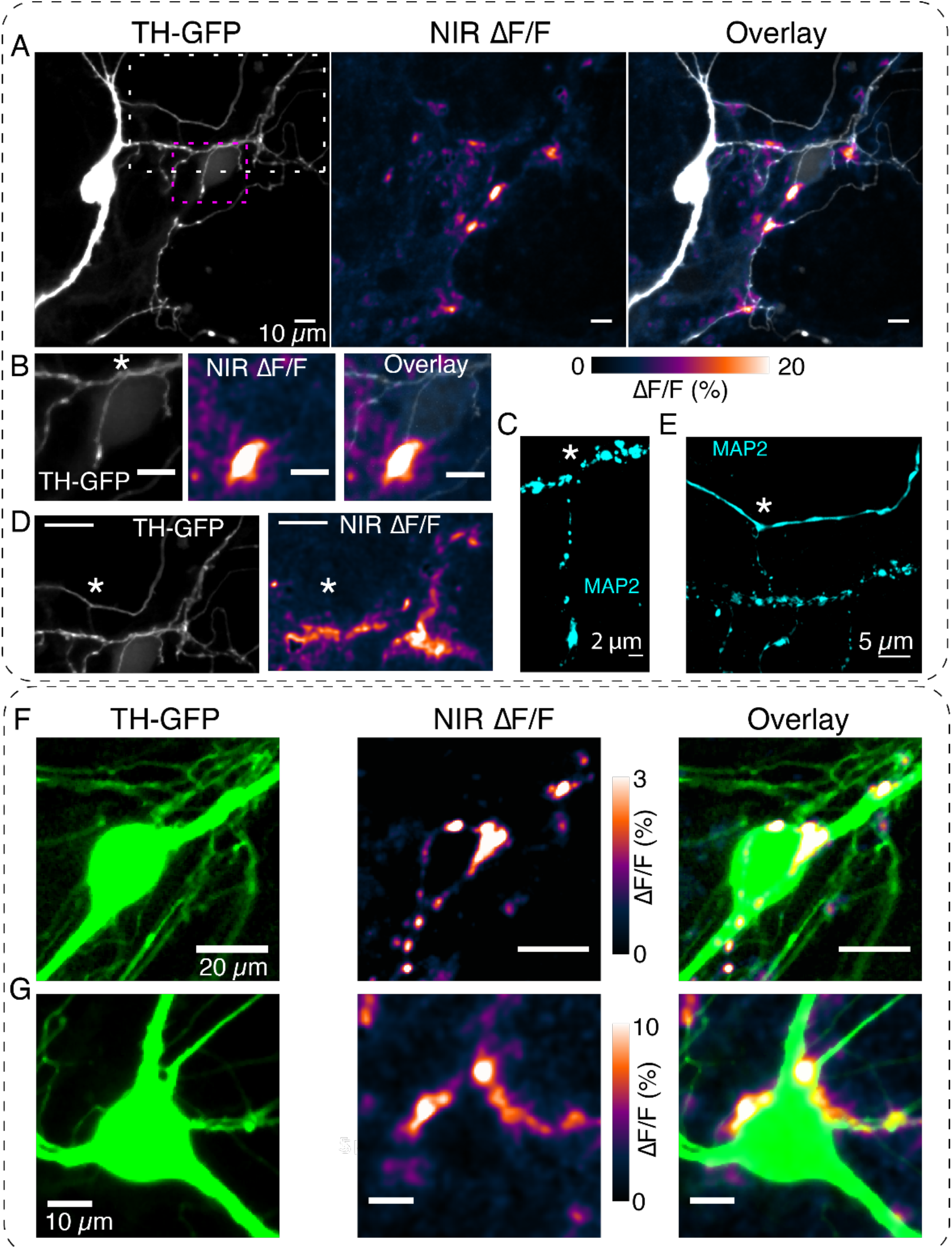
DopaFilm detects dopamine release activity from dendritic processes. (A) TH-GFP image of a dopamine neuron and its spontaneous DopaFilm activity. Maximum intensity projection from a ΔF/F stack is shown, overlayed with TH-GFP. (B) Area bounded by magenta box in (A) and close-up of its spontaneous DopaFilm activity and overlay. Scale bar = 10 µm (C) MAP2 signal corresponding to (B) Use * to compare FOVs in (B) and (C). (D) Area bounded by white box in (A) and close-up of its spontaneous DopaFilm activity. Scale bar = 20 µm. See overlay image in Figure S6. (E) MAP2 signal corresponding to (D). Use * to compare FOVs in (D) and (E). (F, G) DopaFilm activity arising from dendrites that comingle with the soma of dopamine neurons. Notice activity from fine dendritic processes that appear in proximity to the soma. Transient traces for activity in (F) are shown in Figure 2F.

**Figure 6:**
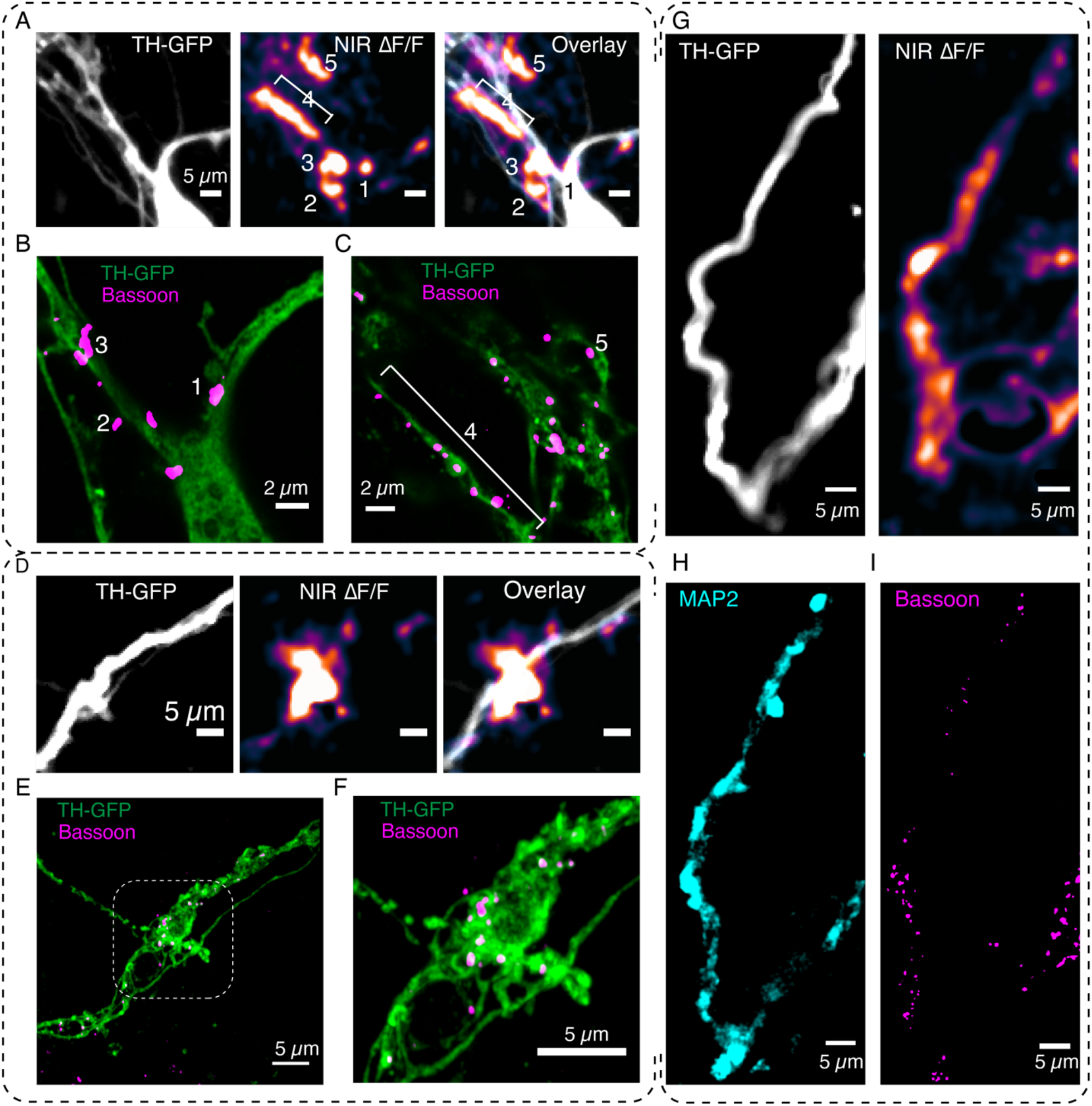
DopaFilm enables interrogation of the molecular correlates of dendritic release. (A) DopaFilm peak ΔF/F activity overlayed with TH-GFP image around soma of dopamine neuron. NIR ΔF/F image contrast set at 0 – 10 % ΔF/F. (B, C) Airyscan super resolution images corresponding to FOV in (A). (D) Dendrite of dopamine neuron and its Δ/F activity. NIR ΔF/F image contrast set at 0 – 10 % ΔF/F. (E) Airyscan super resolution images corresponding to FOV in (D). (F) Close up of boxed region depicted in (E). (G-I) Activity from a dendritic process and its corresponding TH-GFP, MAP2 and Bassoon images. NIR ΔF/F Image contrast set at 0 – 30 % ΔF/F.

Despite the noted differences between axonal and dendritic release, our experiments showed that DopaFilm transients at dendritic processes shared many key features with those in axons. Fluorescence turn-on was fast for both evoked and spontaneous activities (Figure S7, Figure S8, Figure S11, Figure S12), suggesting availability of fusion-ready docked vesicles reminiscent of vesicular release at axonal terminals. Moreover, dendrites could sustain high levels of activity in extended imaging experiments, suggesting an available pool of vesicles that can be quickly recruited, docked, and primed (Figure S11). These data suggested that release of dopamine in dendrites may be mediated by molecular machinery comparable to those in axon terminals.

Moreover, we explored to what extent, if any, somata of dopamine neurons contribute to somatodendritic release. As was often the case in most imaging sessions, we saw little direct activity arising from soma of dopamine neurons (Figure 5A). When DopaFilm activity was noted near soma, such activity often fell into one of two categories. First, we noticed that DopaFilm activity can often be observed on the major dendrites of dopamine neurons (that is, the stereotypic thick dendritic trunks that depart from the soma), including at the junction between the soma and dendrite (Figure S7, Figure S12). Second, we visualized release from dendrites that commingled with the cell body, often coming in such proximity that it appears as if activity arose from the soma (Figure 5F, 5G, Figure S7, Movie S6). However, the subcellular imaging capability of DopaFilm clarifies, even in such cases, that activity was likely driven by dendrites that appeared to innervate the soma.

### DopaFilm enables interrogation of the molecular correlates of dopamine release

We next used DopaFilm to examine the protein machinery involved in organizing release in dendritic processes. In classical synapses, release of neurotransmitters from axon terminals occurs at a highly specialized structure called the synaptic active zone. The active zone is enriched in large scaffolding proteins (Bassoon, Piccolo) and protein complexes that dock and prime synaptic vesicles (RIM, RIM-BP, ELKS, MUNC-13) as well as SNARE-complex proteins (VAMP, SNAP 25 and syntaxin) that carry out vesicle fusion in the final exocytotic step.^2^ We wanted to know if DopaFilm dendritic activity was co-located with the expression of proteins that are classically involved in neurochemical release at axon terminals. To explore this question, we set our sights on Bassoon, a large scaffolding protein that is classically used as a marker for presynaptic terminals in axons,^37^ and whose disruption leads to loss of synaptic transmission.^38^ We first performed DopaFilm activity imaging in dendritic processes of dopamine neurons. We then immunostained for Bassoon and performed post hoc Airyscan imaging at dendritic locations that exhibited hotspots of DopaFilm activity. These experiments showed that Bassoon was enriched at dendritic locations where we observe DopaFilm ΔF/F hotspot activity (Figure 6, Figure S7, Figure S8). In some images, DopaFilm activity hotspots were localized directly to a Bassoon punctum (Figure 6B, 6C, Figure S7), while in others, we observed that DopaFilm activity colocalized with enriched clusters of Bassoon puncta (Figure 6E, 6F). Additionally, when DopaFilm activity was observed along a whole segment of a dendritic process (as opposed to a localized hotspot on the process), we noted that Bassoon is likewise enriched along the process (Figure 6G, 6H, 6I). Importantly, while we observed that presence of Bassoon did not necessarily indicate presence of DopaFilm activity, we did not observe DopaFilm activity from a dendritic process that did not have Bassoon puncta (Figure S12A-D). These results suggest that Bassoon likely plays a key role in organizing dendritic release of dopamine in a manner that is reminiscent of its function in nerve terminals.

We similarly explored if DopaFilm activity co-localizes with proteins involved in the exocytotic release of neurotransmitters. A myriad of cellular functions that require membrane fusion, including subcellular compartmentalization, cell growth and chemical secretion, involve the use of SNARE complex proteins.^39–41^ SNARE proteins are responsible for triggering synaptic vesicle fusion as the last step of chemical neurotransmission.^42^ Previous immunofluorescence studies have established that that midbrain dopamine neurons likely use the vesicular SNARE protein synaptobrevin-2 (also known as VAMP2) as opposed to the more conventional variant synaptobrevin-1 (VAMP1) for exocytotic release.^21, 43^ We explored if the expression of synaptobrevin-2 correlated with DopaFilm activity along a dendritic process. When imaging in dendritic arbors, we observed that DopaFilm activity along a dendritic process exhibited a strong correlation with the expression level of synaptobrevin-2, while TH+ segments with no synaptobrevin-2 expression lacked a corresponding DopaFilm activity (Figure 7).

**Figure 7:**
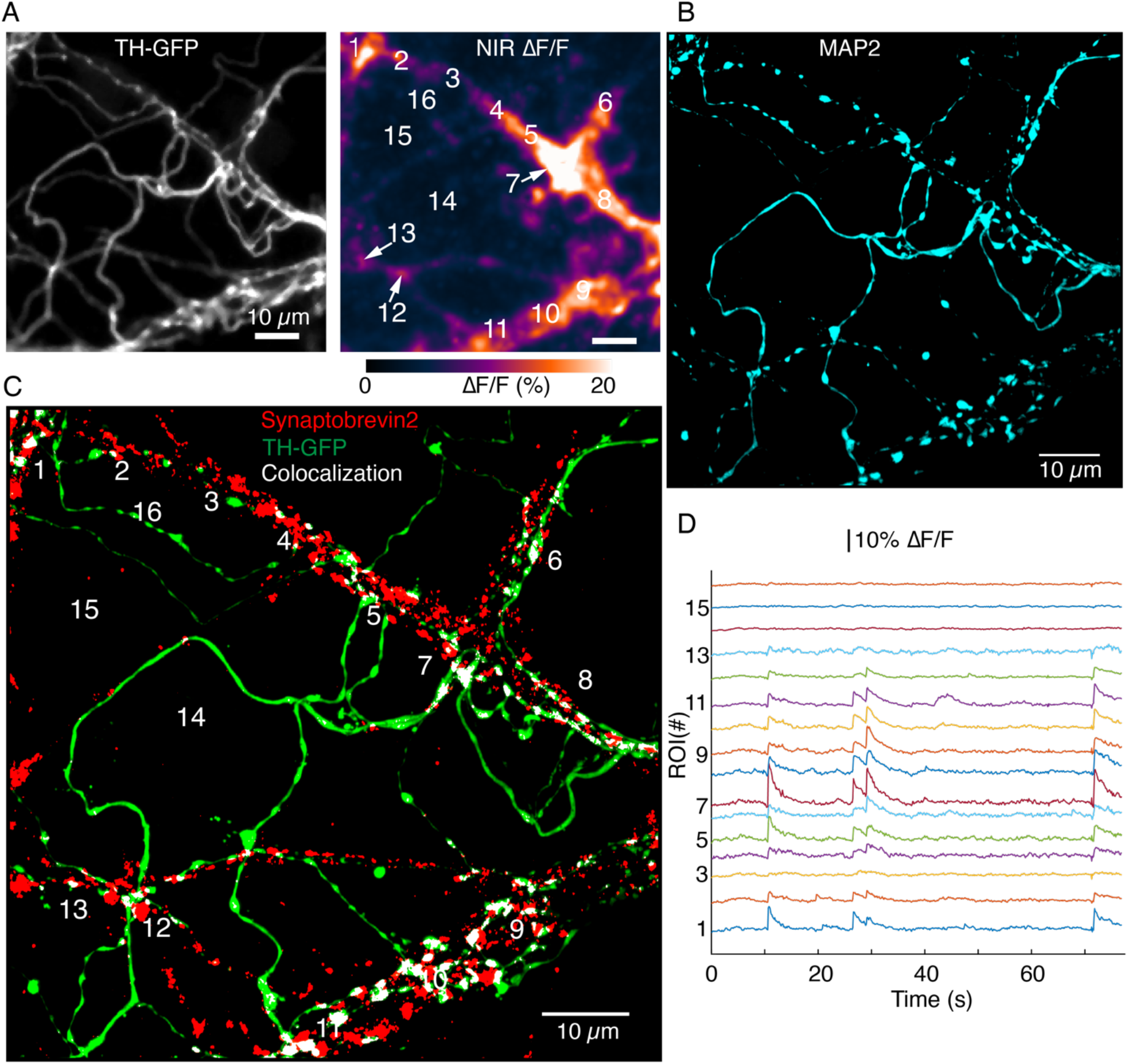
Synaptobrevin-2 is enriched at dendritic locations with high DopaFilm activity. (A) DopaFilm activity image in a dendritic arbor of autonomously spiking dopamine neuron (i.e., no external stimuli applied). Soma not shown. (B) MAP2 Airyscan image of TH-GFP FOV in (A) shows the dendritic nature of processes in (A). (C) Airyscan image of TH-GFP image in (A) showing double-staining for TH-GFP (green) and Synaptobrevin-2 (red). Colocalized green and red puncta are color-thresholded to display white. (D) ΔF/F traces of ROIs 1 to 15 depicted in (A) and (C). Notice that segments that have no synaptorevin-2 signal exhibit no DopaFilm activity (examples: ROI 3, 14, 15, 16).

## Discussion

Among design criteria for biological sensors is the requirement that activity be read out in temporal and spatial domains at sufficiently high sampling rates to recapitulate the underlying dynamics. From a temporal perspective, designing probes with suitable kinetics is carried out routinely during biosensor development. However, targeting the designed probes to read out the biochemical processes at appropriate spatial scales is not always trivial. As it pertains to chemical synapses, the nature of chemical effluxes demands uniform delivery of probes to the synapse, and peri-and extra-synaptic spaces because such signals are inherently incompatible with cell-centered experimental paradigms. In this study, we introduced DopaFilm as a technology that facilitates visualization of chemical effluxes from dopamine neurons. DopaFilm is inherently omnipresent (two-dimensional) and is therefore capable of detecting chemical efflux at the source (synapse) and track the signal to the end of its inherent spatial extent, subject to its limit of detection. We therefore circumvent the non-trivial demand of targeting the probe to the synapse and its surroundings. This design approach, coupled with the film’s favorable temporal properties and quantal sensitivity, has facilitated visualization of synaptic dopamine chemical effluxes with unprecedented resolution. We leveraged these capabilities to shed light on the nature of dopaminergic chemical synapses. We demonstrate that dopamine varicosities serve as beacons that broadcast dopamine release into neighboring peri and extra-synaptic spaces. DopaFilm registered these release sites as hotspots of fluorescence activity and further afforded tracking of the evolution of the hotspots in the spatial domain, in real time.

To demonstrate the utility of DopaFilm, we applied it to study release in somatodendritic compartments of dopamine neurons. Dopamine release in midbrain regions, particularly in the substantia nigra pars compact (SNc) and ventral tegmental area (VTA), are a critical component of dopamine signaling for two important reasons. First, dopamine released in SNc and VTA is predicted to activate D2-autoreceptors, which in turn tune the excitability of dopamine neurons through G protein-gated inwardly rectifying K^+^ (GIRK) channels.^44, 45^ This self-inhibition of dopamine firing in turn regulates dopamine release in distal regions that receive dense axonal innervation from midbrain dopamine neurons, including the striatum and prefrontal cortex, in addition to regulating release in SNc and VTA. Second, dendritic projections from the VTA into the substantia nigra pars reticulata (SNr) are thought to activate D1-receptors and regulate output from the primary GABAergic neurons of the SNr.^21, 46–48^ Through these two mechanisms, somatodendritic release plays a critical role in the wide range of functions that have been ascribed to dopamine neuromodulation, including motor control, motivation, and learning.^21, 45^ Moreover, according to recent studies, somatodendritic release is posited to be responsible for delaying the onset of symptoms in Parkinson’s disease, compensating for the extensive axonal degeneration that SNc dopamine neurons exhibit.^49^ In sum, somatodendritic dopamine release is an important facet of dopaminergic neuromodulation, and plays important roles in health and disease.

Despite this importance however, insights into the spatiotemporal dynamics and regulatory mechanisms of somatodendritic release have been lacking, largely owing to the intermingling between soma and dendrites, the inability of available assays to tease out release at sub-cellular levels and the non-canonical nature of non-axonal release. We therefore deployed DopaFilm to study release in somatodendritic compartments of dopamine neurons. Our results show that somatodendritic release of dopamine arises primarily from the dendrites and not the soma. Major dendrites of dopamine neurons (that is, the thick, smooth fluorescent trunks that emerge from the cell body, before ramification), as well as fine dendritic processes and dendritic arbors participate in dopamine release. Remarkably, dendrites with no apparent varicose morphologies participated in dopamine release as did dendritic arbors with axon terminal-like bouton structures. Dendritic activity shared a lot in common with dynamics observed in axons, but dendritic hotspots appeared localized in their spatial extent to the immediate vicinity of the release site in contrast to release in axonal processes, which propagated to a larger spatial extent. Dendritic release was observed to be robust, Ca^2+^-dependent, and is broadcast from hot-spots that are enriched with the presynaptic active zone protein Bassoon. Our study ascribes an important role for the SNARE-complex protein synaptobrevin-2 in dendritic dopamine release, consistent with predications from previous studies.^43^ Establishing the full complement of necessary and sufficient molecular machinery that organizes dendritic and axonal release would require a systematic experimentation that will be the subject of future studies.

### Context

Although a rapidly growing suite of tools are being developed to facilitate measurement of neurochemical release, we believe that DopaFilm and other similar technologies will fill an important niche and address a known need in neuroscience research. Most genetically encoded biosensors for neurochemicals are primarily optimized for large scale multicellular imaging or fiber photometry experiments.^50–52^ In such applications, biosensors are broadly expressed on the plasma membrane of neurons, often in non-cell specific manner and are not targeted to synapses for single release site assay. Recent studies have sought to redeploy these biosensors to measure presynaptic release, but these approaches still rely on cell-surface anchoring strategies that limit their read out in spatial domain, and it remains to be established whether the sensors are truly presynaptically localized.^53, 54^ Electrical or Ca^2+^ recordings from post-synaptic processes can help us study presynaptic chemical synapses but these methods are lacking in throughput, represent summed synaptic input from multiple release sites and are not suitable for neuromodulators that signal through G-protein coupled receptors.^55–59^ FM dyes, pHluorins and other pH-sensitive vesicle-loadable optical tracers of synaptic vesicles require intense stimulation to drive the endo-exocytotic cycle and are not designed to visualize neurochemical diffusion at the requisite spatial scales.^31, 60, 61^ Nevertheless, carefully designed experiments using pHluorins have enabled studies of presynaptic regulation of chemical synapses.^62–65^ Redox active molecules like dopamine have benefited from amperometric measurements, which have reported quantal release events with sub-millisecond temporal resolution.^66, 67^ Furthermore, GIRK-mediated post-synaptic currents have been leveraged to measure quanta of dopamine release.^68^ Here too, however, measurements are low throughput single point assays, and cannot to be localized to presynaptic release structures, and do not convey spatial information. In conclusion, measurements of neurochemical release with full spatiotemporal resolution and quantal sensitivity have remained elusive. This hampers our ability to visualize and understand synaptic chemical effluxes, to study the heterogeneity of chemical synapses within the same cell or across cell types, to investigate the complex regulatory mechanisms of presynaptic release, and how these get shaped during synaptic plasticity, development, and disease. Therefore, new technologies that augment the capabilities of existing tools can make meaningful contributions to the study of neuroscience.

### Limitations

Synthetic strategies such as DopaFilm can fill a niche in the suite of tools that are designed to study cellular chemical release. Moreover, DopaFilm’s non-photobleaching fluorescence in the NIR to SWIR regions of the spectrum (0.85 to 1.35 µm) is spectrally compatible with existing optical technologies, as we have demonstrated in this study. However, the unique spectral properties of DopaFilm means that microscopes that typically rely on silicon technology for photon detection (cameras and PMTs) are not compatible with its use. DopaFilm requires use of detectors with InGaAs sensors for photon detection, and additional optimization of optical components to facilitate transmission of NIR and SWIR photons. This requires an investment by the end user, albeit a modest one. Furthermore, single wall carbon nanotube-based nanosensors, from which DopaFim is fabricated, have been developed for electroactive neurochemicals (catecholamines, indolamines)^22, 69^ but strategies to sense molecules such as glutamate and GABA, and neuromodulators such as neuropeptides are still active areas of research. This would restrict DopaFilm-like strategies to the study of molecules for which we have robust nanosensors now. Finally, while synthetic probes do not have the benefit of ease of use that genetically encoded probes possess, they lend themselves to deployment in unique preparations such as one demonstrated in this study and can offer opportunities to study neurobiology in less genetically accessible regions of the brain and in genetically inaccessible organisms.

## Acknowledgments

This study was supported by the Howard Hughes Medical Institute through Janelia Research Campus (JRC). We thank Kristen Delevich, Joshua Dudman, Markita Landry, Luke Lavis, Timothy Ryan, Eric Schreiter, David Stern, and Linda Wilbrecht for discussions and comments on the manuscript. We express our gratitude to Dmitri Tsyboulski (Janelia Experimental Technology), and Wei Sun and Paulo Chaves (Thorlabs) for assistance with development of the visible-SWIR broad spectrum imaging microscope. We thank the Molecular Biology and Viral Core facilities at JRC for their assistance and vivarium team for support with animal husbandry. We thank Kaspar Podgorski for developing and sharing with us the MATLAB program used for data analysis and David Ackerman for improving the program to suit our needs. We thank Shu-Hsien Sheu and Boaz Mohar for sharing reagents and for helpful discussions on experiments. We are grateful for the support provided by imaging specialist Damien Alcor and the Advanced Imaging Center team at JRC.

## Supplementary Figures

**Figure S1:**
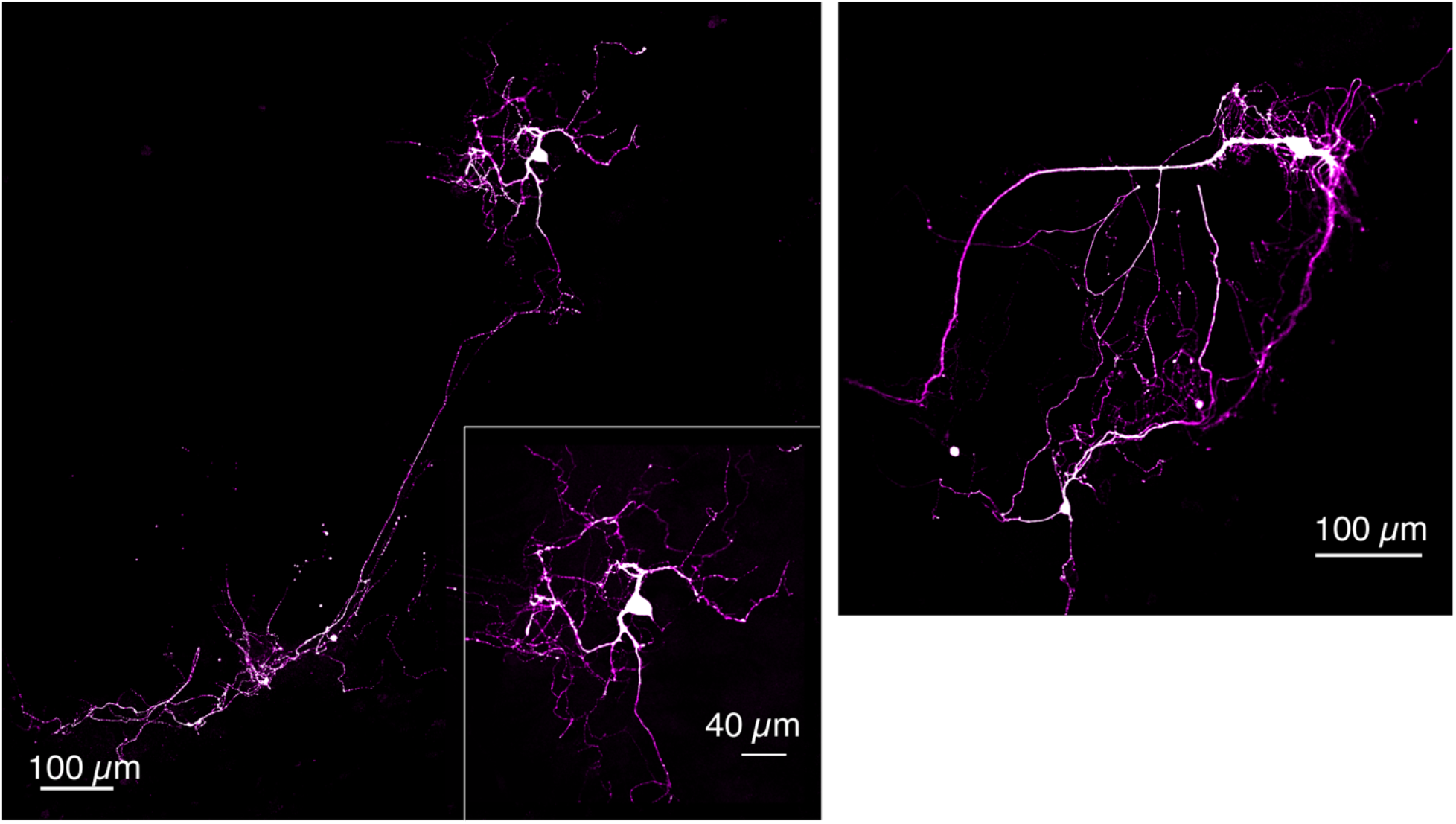
TH-immunofluorescence of Dopamine neurons grown on DopaFilm. Examples of TH+ dopamine neurons grown in co-culture with cortical neurons.

**Figure S2:**
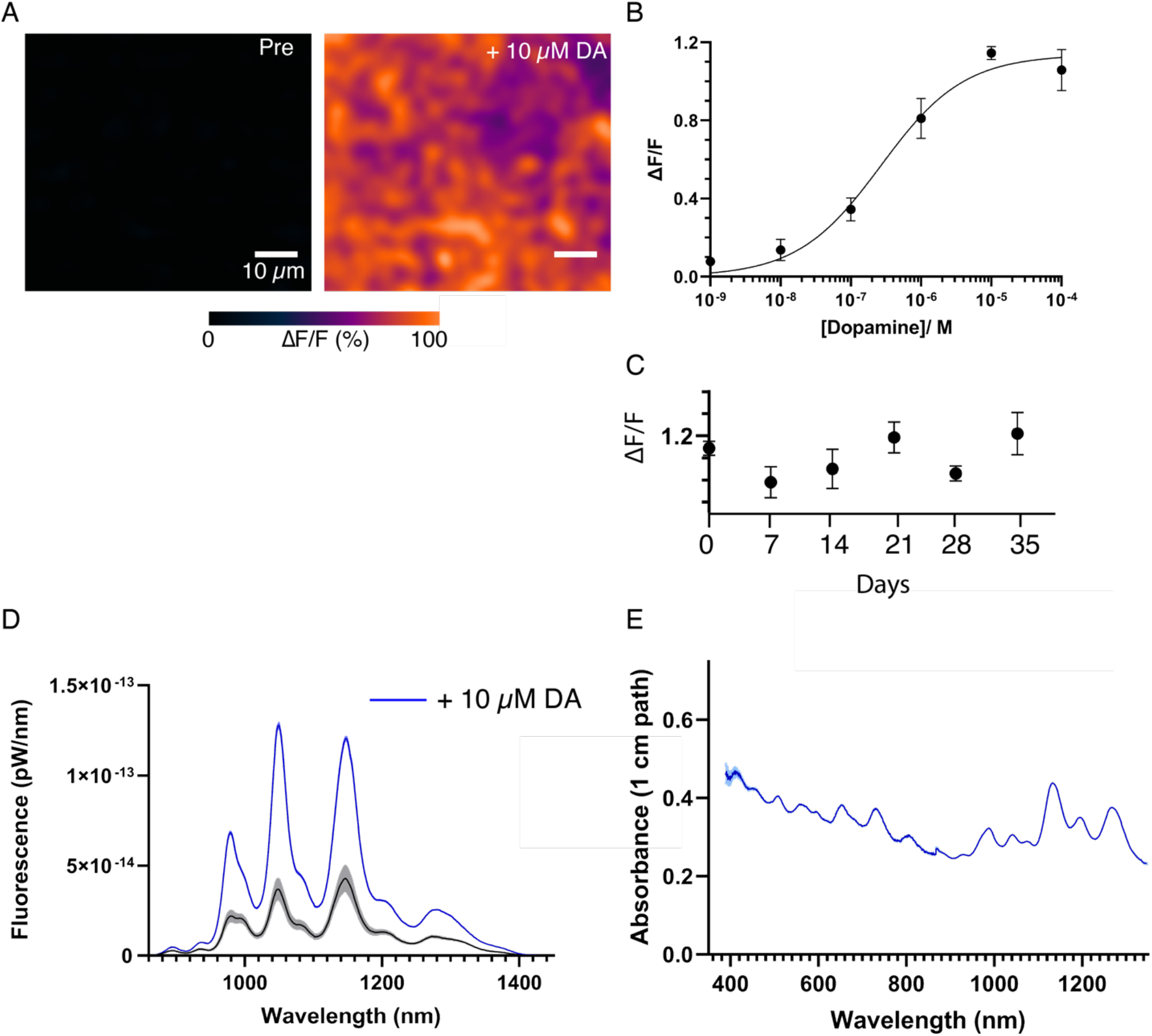
DopaFilm and solution phase nanosensor characterization. (A) 10 µM of exogenous dopamine (DA) wash on DopaFilm shows diffuse ΔF/F response from DopaFilm substrate. (B) Dose-response curve of DopaFilm. Each experimental data point has an error bar depicting standard deviation from different DopaFilm preparations (n = 3). Experimental data points were fit to the Hill equation (least-squares regression, black trace) to estimate an apparent dissociation constant (*K_d_ = 268 nM*) and Hill coefficient (*n = 0.73*). (C) DopaFilm response to 10 µM DA remained stable over 35 days. Each experimental data point has an error bar depicting standard deviation from different DopaFilm preparations (n = 3). x-axis: days post initial preparation. (D) Solution phase fluorescence emission spectra of dopamine nanosensors, prepared from a conjugation of multi-chiral single wall carbon nanotubes functionalized and single strand oligonucleotides, before (black trace) and after (blue trace) application of 10 µM DA. Bands indicate standard deviation from n = 3 measurements. (E) Solution phase absorption spectrum of nanosensors.

**Figure S3:**
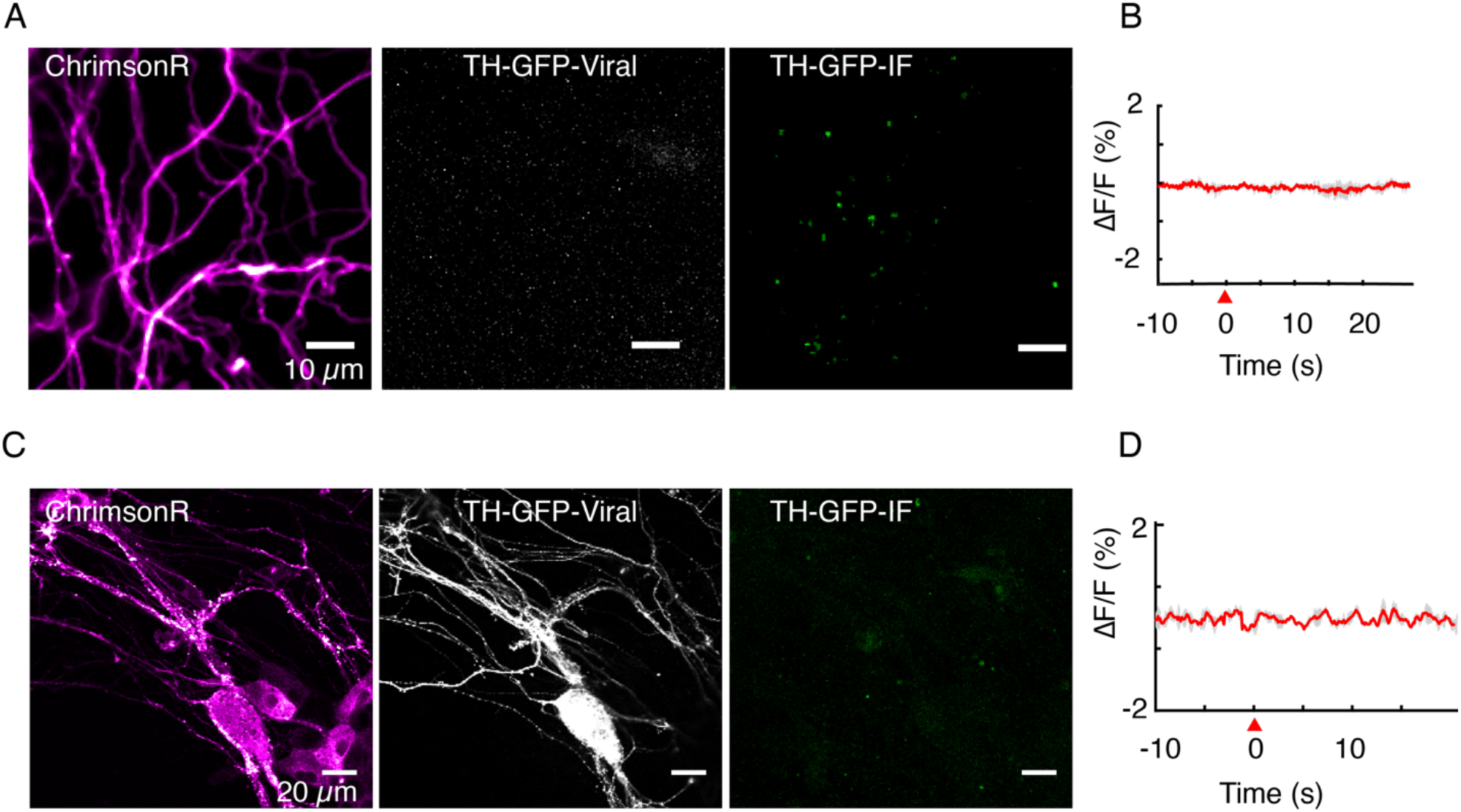
TH immunofluorescence is used to verify identity of putative dopamine neurons. (A, B) Optical stimulation of ChrimsonR-positive and TH-GFP-negative axonal arbor elicits no evoked activity (n = 25 cells). Immunofluorescence (TH-GFP-IF) indicated no dopaminergic processes in the field of view. (C, D) Optical stimulation of ChrimsonR-positive and TH-GFP-positive soma and dendritic arbor elicits no evoked activity (n = 10 cells). Retrospective immunofluorescence (TH-GFP-IF) indicates there are no dopaminergic processes in the field of view. Red wedge: time of application of optical stimulation. Data in (B) and (D) are averaged from triplicate stimulations for FOVs shown in (A) and (C) respectively and are not averages across all cells images.

**Figure S4:**
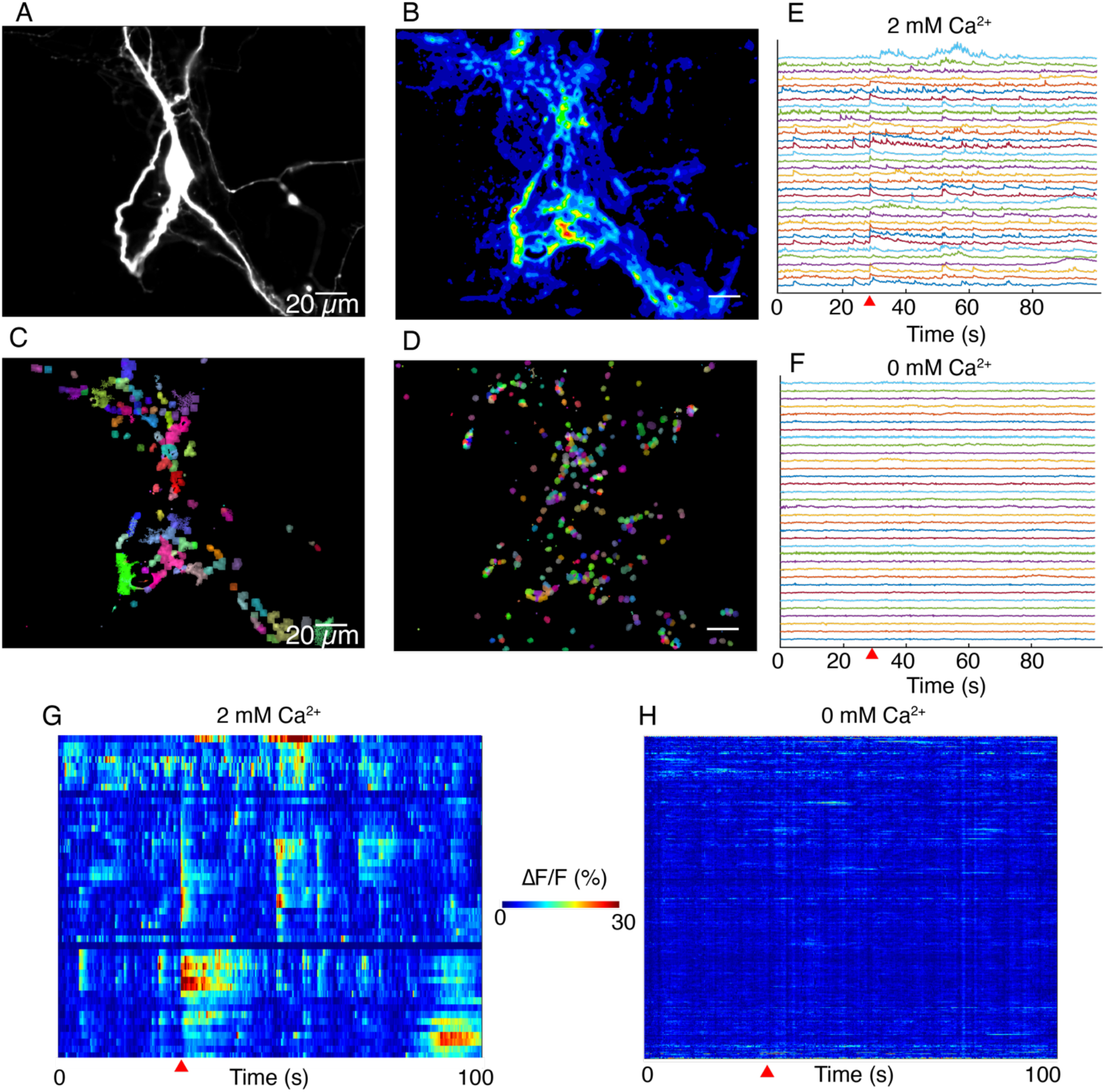
DopaFilm activity imaging in normal imaging buffer (ACSF with 2 mM Ca^2+^) and buffer with no extracellular Ca^2+^ (Ca^2+^-free ACSF). (A) TH-GFP image of a dopamine neuron. (B) Maximum intensity projection from a 1000-frame ΔF/F movie of evoked and spontaneous activity imaging of neuron shown in (A), in normal ACSF. (C) Non-negative matrix factorization (NNMF) decomposition applied to the ΔF/F movie in normal ACSF shows local clusters of pixels with highly correlated activity, color coded for better visualization. Similar colors in a local cluster of pixels corresponds to correlated activity. (D) NNMF decomposition of movie stack of imaging in Ca^2+^-free ACSF. (E, F) ΔF/F time traces for a subset of the NNMF clusters (i.e., decomposed elements) from (C) and (D) respectively, off-set in y-axis for better visualization. (G, H) Heat-maps of activity for all the NNMF components in (C) and (D) respectively. Red wedge in figures (E)-(H) = time of optical stimulus application.

**Figure S5:**
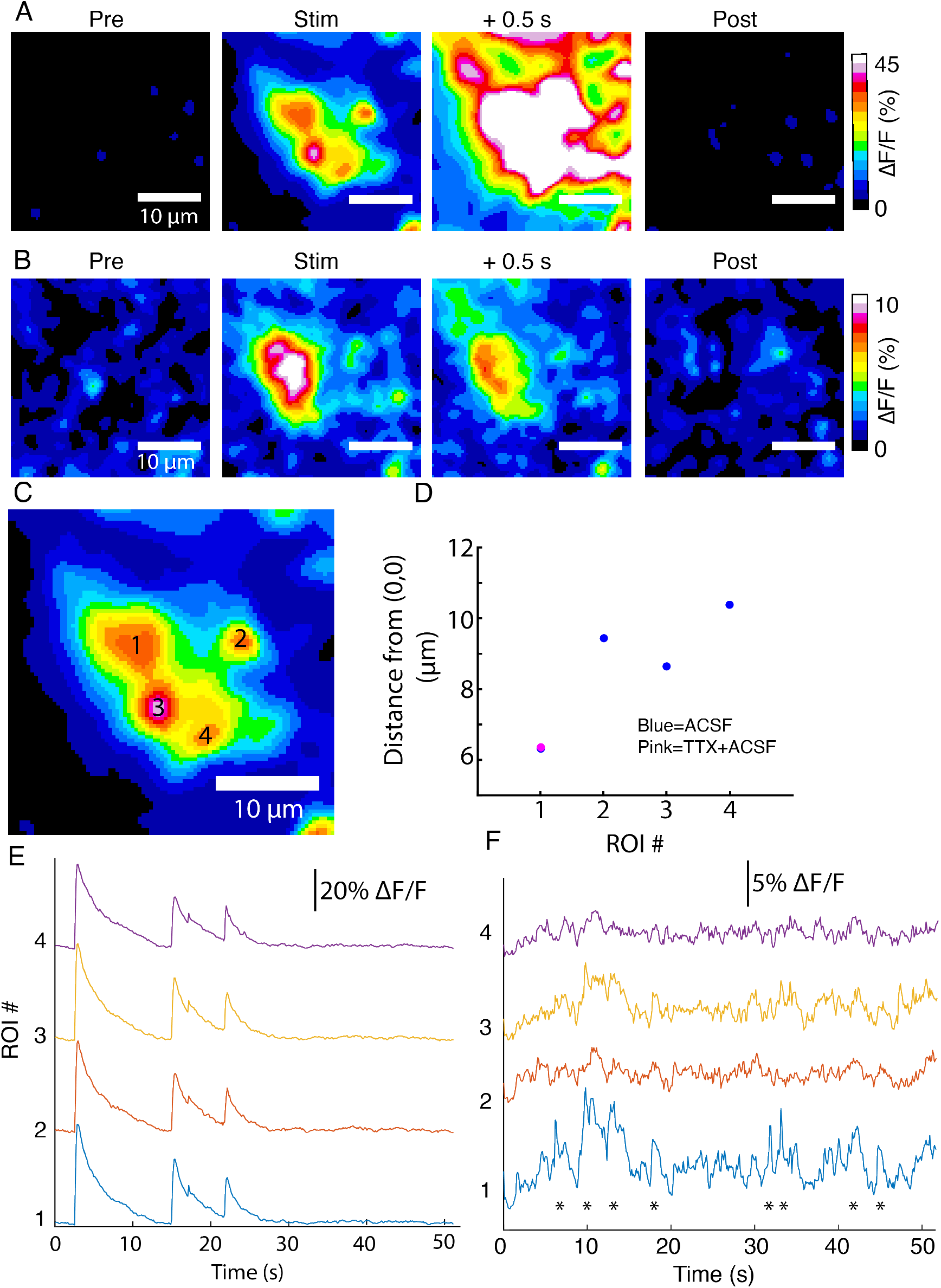
DopaFilm activity imaging in an axonal arbor in: (A) normal ACSF. (B) ACSF with 10 µM of TTX. (C) DopaFilm activity is driven by four putative release sites when imaging in ACSF, but activity is only seen in ROI #1 when TTX is applied. (D) Centroid of hotspots in ACSF and ACSF+TTX. For ROI 1, centroids nearly overlap. For ROI2, 3 and 4, TTX centroids are missing. (E, F) ΔF/F activity traces for ROIs 1 to 4 in ACSF (E) and ACSF+TTX (F).

**Figure S6:**
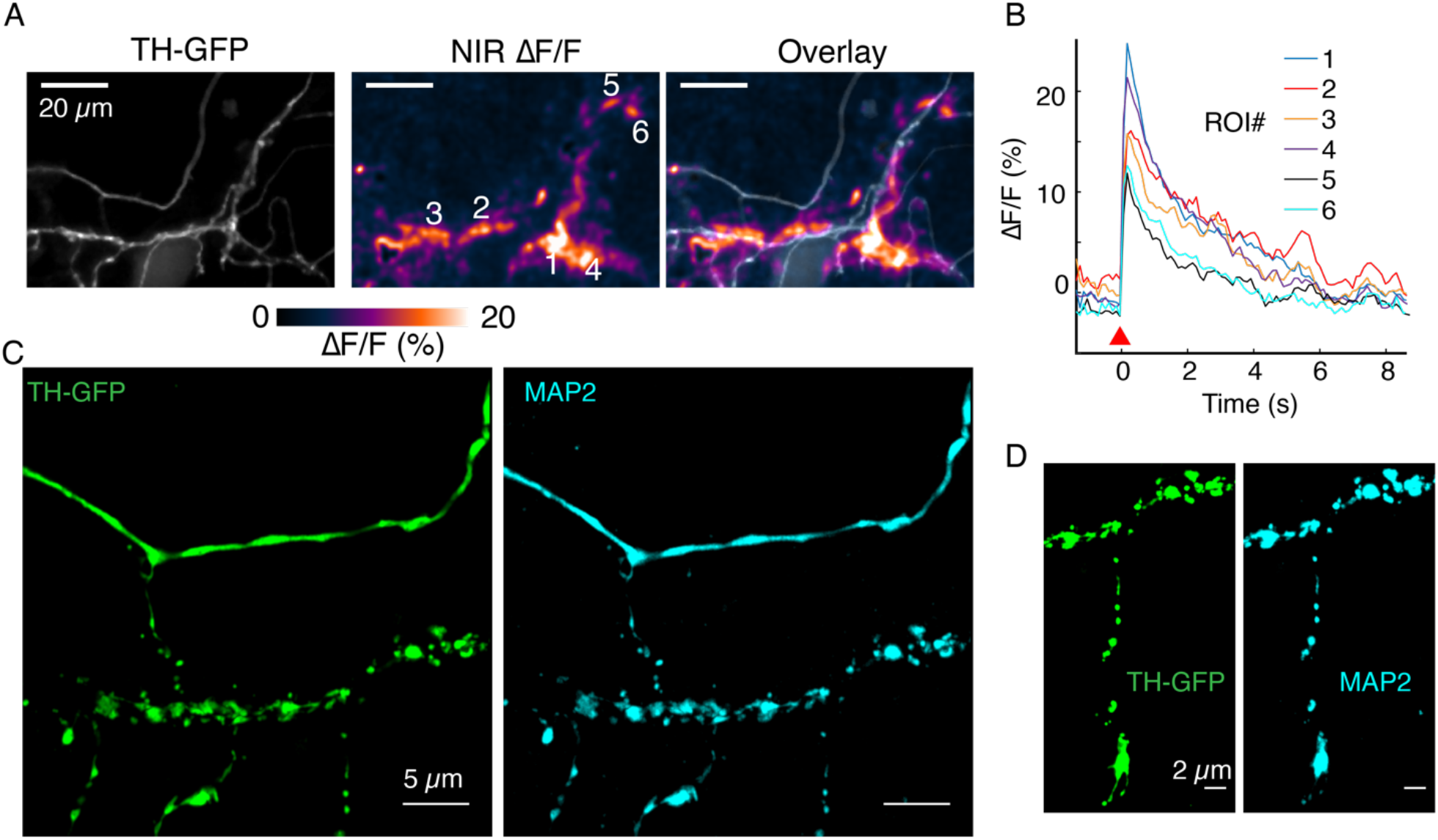
MAP2+ immunofluorescence demonstrates dendritic nature of processes. (A) TH-GFP, NIR ΔF/F and overlay image corresponding to data in Figure 5D. (B) Activity traces corresponding to ROIs 1 to 6 noted in (A). (C) TH-GFP and MAP2 double-stain and super resolution image of FOV in (A). (D) TH-GFP and MAP2 double stain of FOV in Figure 5B.

**Figure S7:**
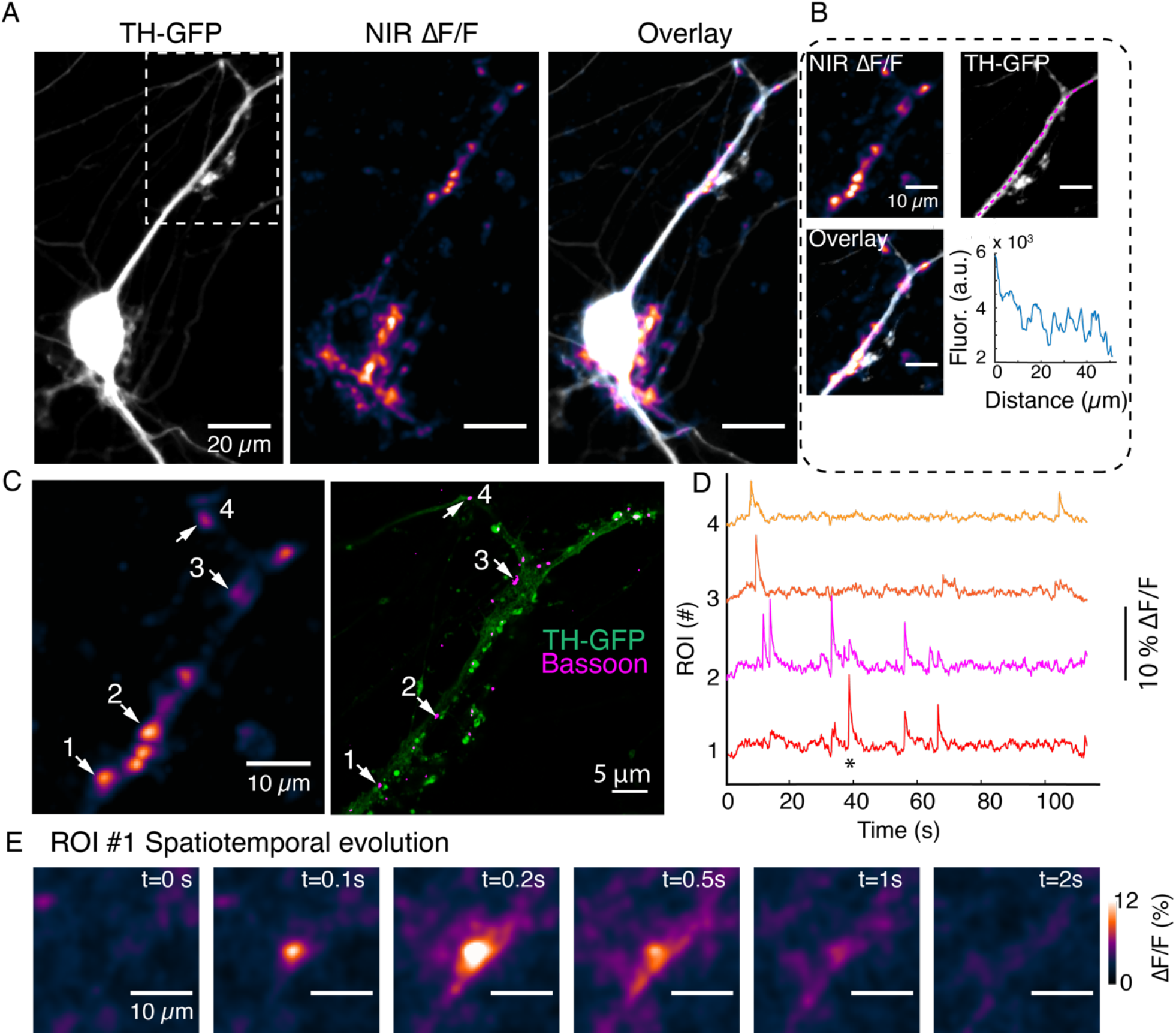
DopaFilm activity imaging at a dendrite of autonomously spiking dopamine neuron. (A) TH-GFP image of a dopamine neuron, and maximum intensity projection from ΔF/F stack and overlay. (B) Segment of activity in (A), (white box) and its TH-GFP intensity profile along the dendritic process (along magenta line), show no putative boutons. (C) DopaFilm hotspot activity for the selection depicted in (A) and corresponding TH-GFP and Bassoon super resolution images. Bassoon puncta can be assigned to DopaFilm hotspot activity. (D) ΔF/F traces for ROIs 1 to 4 shown in (C). (E) Spatial dynamics of ROI#1 at the time depicted with * in the ROI1 trace shown (D)

**Figure S8:**
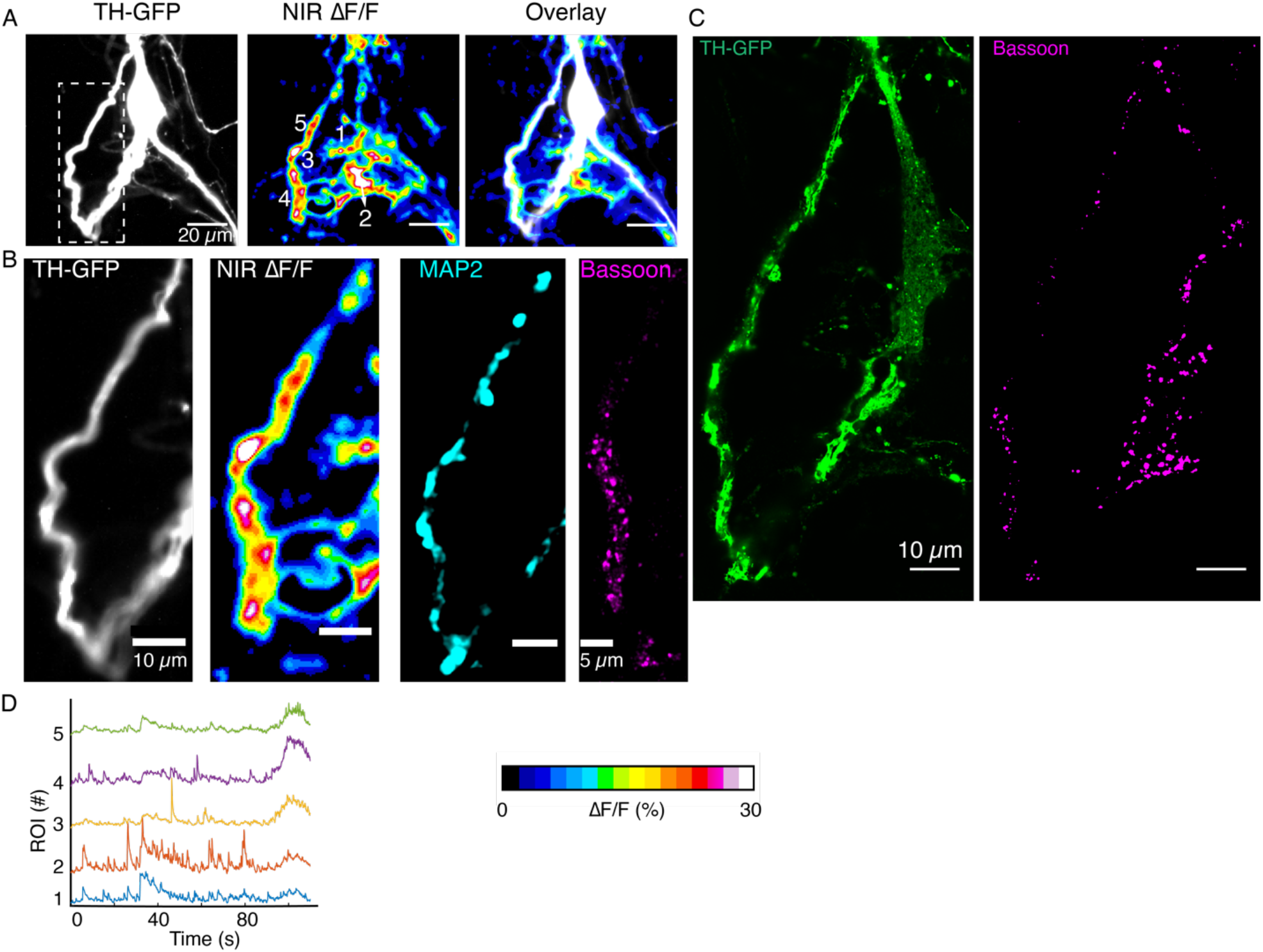
DopaFilm activity imaging of a spontaneously active dopamine neuron. (A) TH-GFP image of a dopamine neuron, and maximum intensity projection from ΔF/F stack and overlay (B) TH-GFP for the white box depicted in (A) and its DopaFilm activity along the dendritic process, as well as MAP2 and Bassoon super-resolution images. (C) TH-GFP and Bassoon super-resolution images for the bigger field of view in (A). (D) ΔF/F time traces for ROIs 1 to 5 depicted in NIR ΔF/F panel in (A).

**Figure S9.**
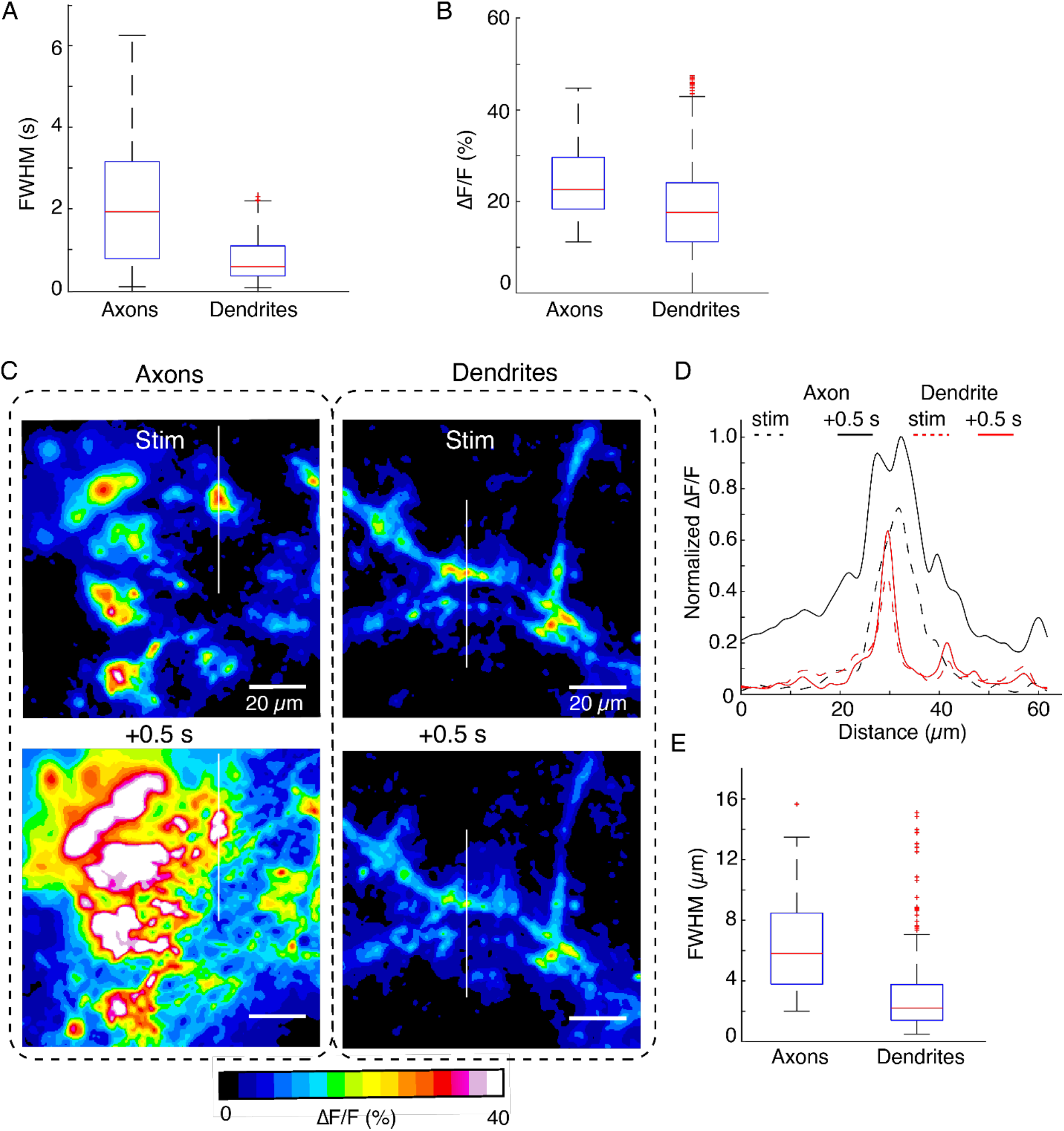
Comparison of spatiotemporal dynamics of DopaFilm hotspots in axonal arbors and dendritic processes after optical stimulation (evoked). (A, B) Box plots comparing pooled data for temporal full width at half-max (FWHM) (s) of transient traces and peak ΔF/F in axons and dendrites. Here, FWHM is defined in the time domain as a measure of how long (in seconds) the transients persist before diffusive and dopamine transporter (DAT)-mediated clearance. Mean ± SD: Peak ΔF/F (%) in axons: 25 ± 8.3. In dendrites: 18 ± 9.3. Unpaired t-test: *p* < 10^-4^. Temporal FWHM (s) in axons: 1.75 ± 1.2. In dendrites: 0.9 ± 0.7. Unpaired t-test: *p* < 10^-4^. (C) Spatial spread of DopaFilm activity in an axonal arbor (left column) and dendritic arbor (right column). ΔF/F images are shown immediately following stimulation (stim, top row) and 0.5 seconds after stimulation (+0.5 s, bottom row). Notice the diffusive broadening of the signal in the axons, which occurs only to a limited extent in dendrites. (D) Line profiles along the white lines depicted on the images in (C). Dash lines represent profile immediately following stimulation (stim) and solid lines represent line profiles after 0.5 s has elapsed. ΔF/F is normalized to the peak ΔF/F observed in the axonal arbor. Notice how activity in axons has a bigger spatial extent than those in dendrites. (E) Spatial FWHM (µm) in axons vs. dendrites. Mean ± SD; axons: 6.6 µm ± 3.6 µm; dendrites: 3.2 µm ± 3µm. Unpaired t-test: *p* < 10^-4^. Data is from n = 32 dendritic processes pooled from n = 12 dopamine neurons. For axons, we pooled data from n = 6 axonal arbors. Boxplot definitions: red line = median, edges of box: 25^th^ and 75^th^ percentile, top and bottom hash lines: minimum and maximum values of non-outlier data, red points: outlier data.

**Figure S10:**
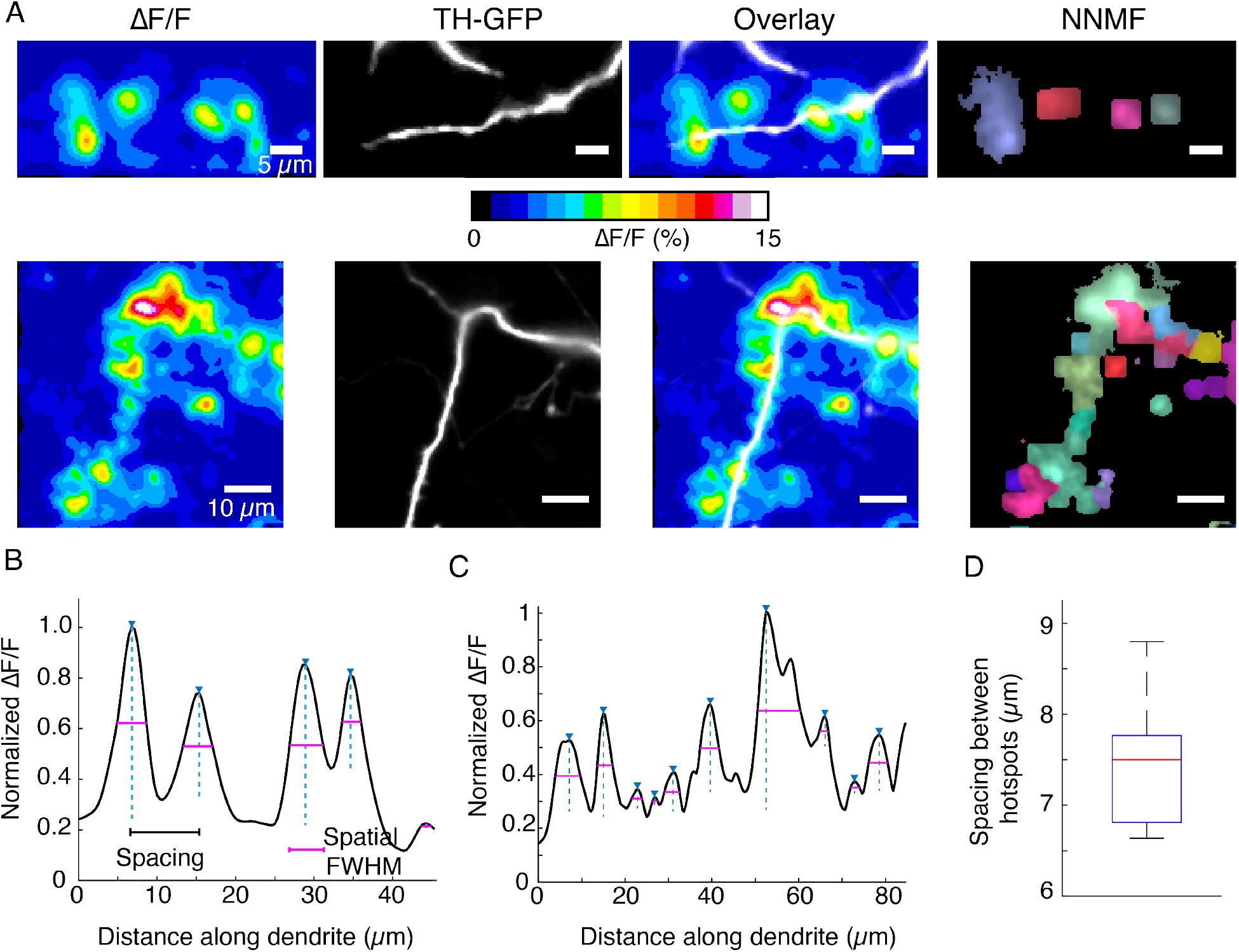
DopaFilm hotspot activity along dendritic processes. (A) Top and bottom rows: ΔF/F, TH-GFP and overlay. Right-most panels: NNMF decomposition of ΔF/F stack identifies clusters of pixels with highly correlated activity, consistent with the ΔF/F hotspots. (B, C) Line profiles of ΔF/F computed along the profile of the dendrites corresponding to the top and bottom rows in (A), respectively. Custom code identifies peaks, distance between peaks (spacing, black line segment) and the spatial extent of each signal, approximated by the FWHM (magenta line segment). (D) Box plot of spacing between hotspots observed on dendritic process. Box plot distribution for FWHM (µm) is shown in Figure S9. Data is pooled from n = 32 dendritic processes pooled from n = 12 dopamine neurons. Boxplot definitions: red line = median, edges of box: 25^th^ and 75^th^ percentile, top and bottom hash lines: minimum and maximum values of non-outlier data.

**Figure S11:**
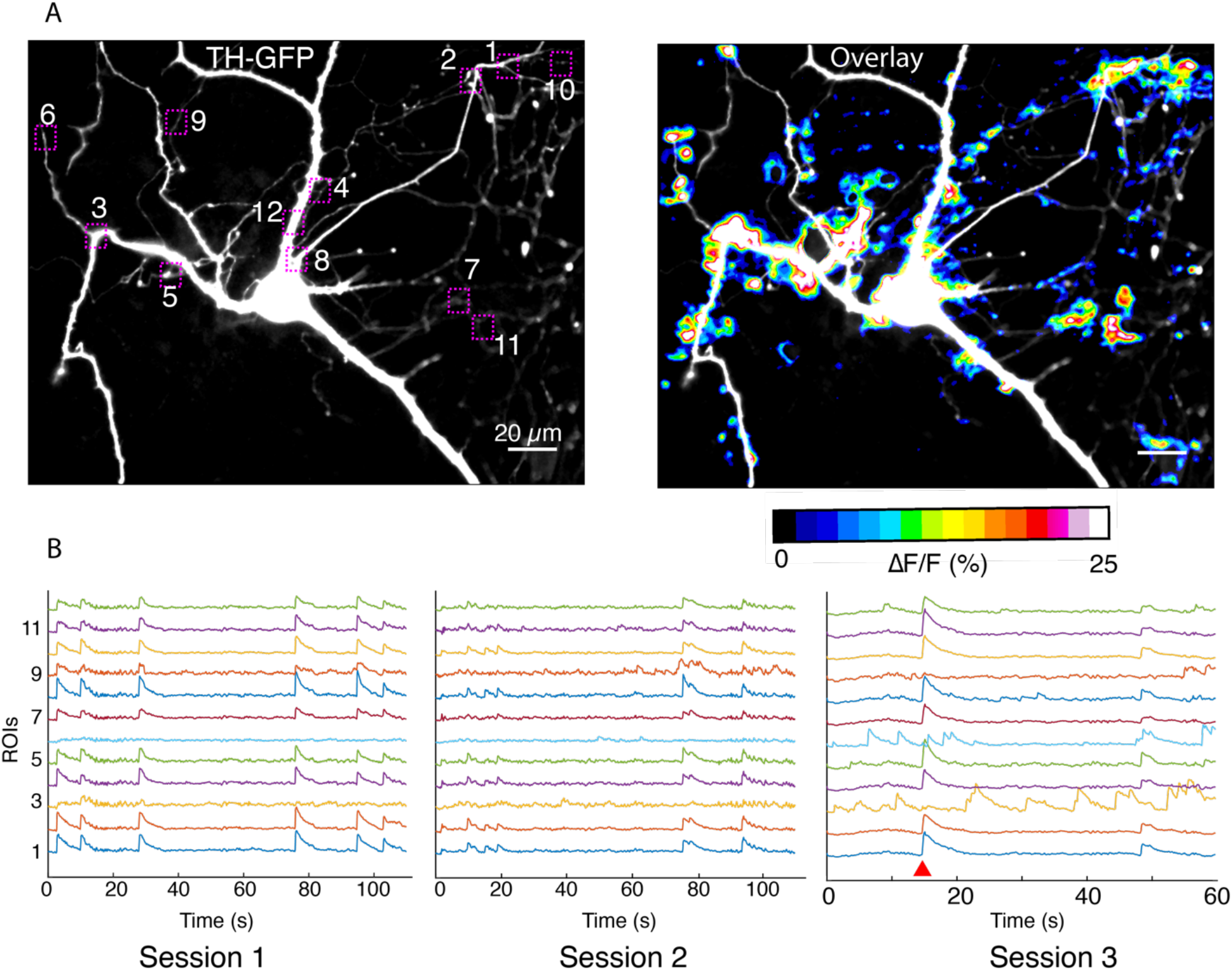
Imaging in dendritic processes of a dopamine neuron. (A) TH-GFP image and its overlay with evoked ΔF/F DopaFilm activity image. (B) ΔF/F traces for the ROI boxes numbered in (A) from n = 5 imaging session (n = 3 sessions shown here). Note that most activity is driven by autonomous spiking except where the red wedge indicates optical stimulus driven activity (in session #3).

**Figure S12:**
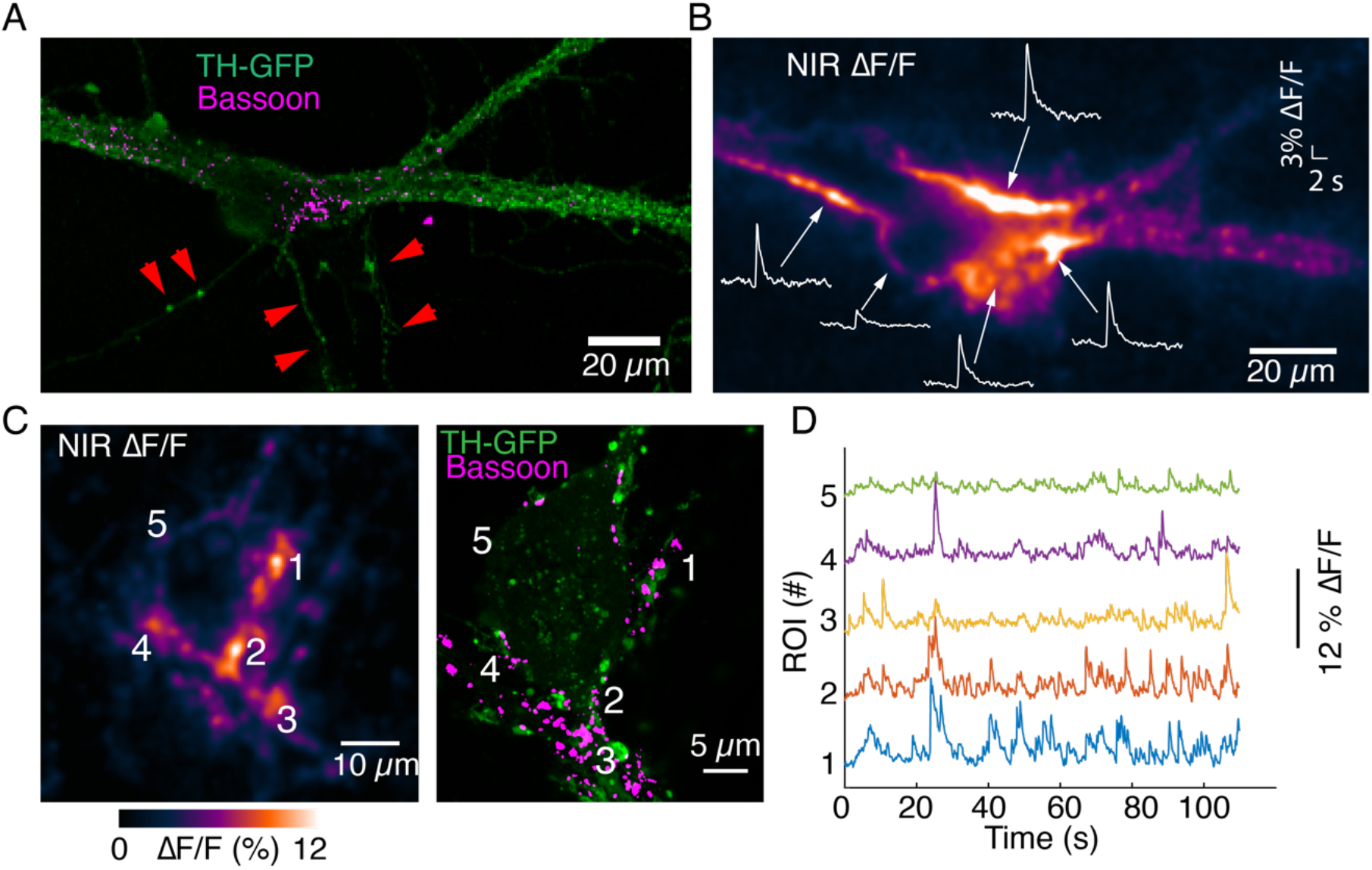
Imaging activity from proximal dendrites associated with the soma of dopamine neurons. (A) TH-GFP image and Bassoon puncta on the major dendrites of a dopamine neuron. (B) ΔF/F activity evoked by optical stimulation. Notice activity arising from major dendritic trunks and at the junction of soma and major dendrites. Also notice lack of activity in processes that lack Bassoon puncta (red arrows). (C) Autonomous spiking activity around the soma of a dopamine neuron. Max ΔF/F projection is shown on left. Notice activity at the major dendrite leaving the cell body (around area depicted as ROI #3) and comingling dendrites (around area depicted as ROI #1, ROI#4), both enriched in Bassoon puncta. ROI#5 exhibits reduced activity relative to ROIs #1 to #4 and has lower levels of Bassoon expression. (D) Activity traces corresponding to ROIs depicted in (C).

**Movie S1:** Imaging in axons.

**Movie S2:** Imaging in axons before application of TTX.

**Movie S3:** Imaging in axons after application of TTX.

**Movie S4:** Evoked release from a dendritic process.

**Movie S5:** Spontaneous activity from a dendritic process.

**Movie S6:** Dendritic activity around soma of a dopamine neuron.

## Methods

### Nanosensor synthesis and characterization

Single-stranded 5′-GTG TGT GTG TGT-3′ [(GT)_6_] DNA oligonucleotides were purchased from Integrated DNA technologies (standard desalting, lyophilized powder) and HiPco single wall carbon nanotubes were purchased from Nano Integris (batch # HR35-141). Solution phase nanosensor synthesis was carried out by first mixing 1 mg of (GT)_6_ and 1 mg of HiPco SWNT in 1 mL of 1X PBS. Then the solution was bath-sonicated (Branson 1800) at room temperature for 20 mins followed by probe-tip sonication (Sonics Vibra Cell) for 15 mins in an ice-bath. Resulting suspension was centrifuged at 20,000 rcf (Eppendorf 5430R) at 4°C for 60 mins and supernatant was carefully transferred into a new Eppendorf tube and stored at 4°C until further use. For characterization, each sensor batch was diluted to a working concentration of 5ppm in 1X PBS. Fluorescence, absorbance, and Raman spectroscopies were carried out on NS Super NanoSpectralyzer (Applied Nanotechnologies). Solution phase fluorescence measurements and dopamine response tests were carried out on a custom built near infrared plate reader from 96-well plates.

### DopaFilm fabrication

Fabrication of DopaFilm on glass substrates was initiated by a silane-based surface modification reaction. First, 35 mm gridded dishes (MatTek P35G-1.5-14-C-GRID) were cleaned sequentially with 200-proof ethanol and copious amounts of molecular biology grade water. The cleaned dishes were incubated with 1 mL of 1% (3-aminopropyl) triethoxysilane (APTES; Sigma Aldrich) in ethanol for 1hour at room temperature. After silane functionalization, dishes were rinsed three times with 2.5 mL of molecular biology grade water. Subsequently, 500 µL of 15 ppm nanosensor solution was added to the glass coverslip of the dish dropwise, allowing the nanosensor to spread evenly. After an overnight incubation at room temperature, excess nanosensor solution was aspirated and carefully rinsed with 2.5 mL of molecular biology grade water. To facilitate neuronal growth on DopaFilm, 1 mL of 0.05 mg/mL poly-D-lysine hydrobromide (PDL; Sigma Aldrich P6407) was applied on top and incubated at room temperature for 1 hour. Finally, dishes were thoroughly rinsed three times with 2.5mL of molecular biology grade water and stored in sterile 1X PBS until seeding neurons. Neurons were seeded directly on the engineered surface immediately after aspiration of the storage PBS solution.

### Neuron co-culture on DopaFilm and viral infections

Primary rat neuronal culture work was conducted according to the Institutional Animal Care and Use Committee (IACUC) guidelines of Janelia Research Campus of the Howard Hughes Medical Institute. Neonatal rat pups were euthanized, cortical hemisphere tissue and substantia nigra pars compacta (SNc) tissue dissected and dissociated in papain enzyme (Worthington Biochemicals) in neural dissection solution (10 mM HEPES pH 7.4 in Hanks’ Balance Salt Solution) at 37°C water bath. After 30 mins, enzyme solution was aspirated, and tissue pieces were subjected to trituration in 10% fetal bovine serum containing MEM media. Following trituration, cell suspension was filtered and resulting single cell suspension was centrifuged to yield a cell pellet. Cell pellet was resuspended in Plating media (28 mM glucose, 2.4 mM NaHCO_3_, 100 µg/mL transferrin, 25 µg/mL insulin, 2 mM L-glutamine, 10% fetal bovine serum in MEM) and cell counts were recorded prior to seeding on DopaFilm substrate. To obtain healthy dopaminergic cultures, mid-brain cells were seeded as co-cultures between SNc cells and cortical hemisphere cells at a ratio of 10:1 (∼300k cells per 35 mm dish). Initially, cells were seeded in attachment media [1:1 (*v*/*v*) plating media to NbActiv4 (BrainBits-NB4)] and cultures were maintained in a 5% CO_2_ humid incubator at 37°C to facilitate cell adherence. After ∼2 hours, attachment media was aspirated and growing media consisting of plating media: NbActiv4 at 1:20 ratio (*v*/*v*) was used. After the first week, cultures were fed twice a week by replacing old media 1:1 (*v*/*v*) with fresh NbActiv4 media. To express opsins and identify dopaminergic neurons, cultures were infected at 5 days in vitro (DIV) with titer-matched viruses using pAAV-Syn-ChrimsonR-tdT (Addgene #59171) and pAAV2.5-TH-GFP (Addgene #80336), respectively (10^5^ combined infectious units per 1 mL of neuronal growth media). Plasmid DNA preparations and virus packaging were performed by Molecular Biology and Viral Tools facilities at Janelia Research Campus.

### Microscopy and Imaging

For broad-spectrum (visible to SWIR, 400 nm – 1400 nm) imaging, we developed a custom microscope based on Thorlabs Bergamo microscope body. We modified the Bergamo to facilitate optimal imaging in 400 nm to 1400 nm with widefield epifluorescence and confocal laser scanning modalities, using commercially available or custom ordered optical components that maximize photon collection, transmission, and detection in the 400 nm – 1400 nm range. We used the widefield epifluorescence modality of the microscope for all our imaging experiments in this study. The microscope is equipped with two fiber-coupled NIR lasers to excite far red, NIR and SWIR fluorophores: a 671 nm laser (GEM 671, Laser Quantum) an 785 nm laser (Thorlabs S4FC785). Additionally, the microscope is equipped with the following 4-wavelength LED light source (Thorlabs LED4D067) to excite and/or actuate visible range fluorophores and opsins: 405 nm, 470 nm, 561 nm, 625 nm. The microscope is equipped with an InGaAs camera with optimized sensitivity in the SWIR range (Ninox 640 II, > 85% QE in 1000 nm – 1500 nm, Raptor Photonics) and an sCMOS camera (CS2100M-USB, Thorlabs) for visible range imaging.

Primary mid-brain neuronal cultures were imaged at 35-40 DIV. At the beginning of a typical imaging experiment, we replaced the NbActiv4 media with artificial cerebrospinal fluid (ACSF) composed of, in mM: 124 NaCl, 2.5 KCl, 1.25 NaH_2_PO_4_, 24 NaHCO_3_, 12.5 Glucose, 5 HEPES, 2 CaCl_2_, 2 MgSO_4_, spiked with 5 mg/L of nanosensor solution for 1 hour. We then replaced the media with fresh ACSF and mounted the culture on the microscope for imaging. We used a 10X objective (Nikon N10XW-PF, 0.3NA, 3.5 mm WD) to identify a dopaminergic neuron from the co-culture system. We then switched to 40X objective (Nikon N40X-NIR, 0.8 NA, 3.5 mm WD) to carry our activity imaging in the identified dopamine neuron. Before each imaging session, we record a GFP image of the TH signal in the green channel. We then switch to the NIR/SWIR channel for activity imaging. Imaging is carried out at frame rates of 10 – 20 frames per second depending on the brightness of the DopaFilm substrate (50 – 100 ms. exposure time). If the neuron exhibits autonomous spiking activity, we do not apply any external stimuli. If the neuron does not exhibit spiking activity, we apply optical stimulus to evoke activity. Optical stimulation is driven through a custom-built MATLAB code that simultaneously controls and synchronizes the camera and optogenetic stimulation light source through external TTL pulses and time-locks stimulation times with specific camera frames. Optical stimuli applied to evoke activity are: 5 pulses, 25 Hz, at power of 1 mW / mm^2^. We use ChrimsonR to evoke activity and use 561 nm LED (Thorlabs LED4D067 with DC4100 driver) of the four-wavelength LED light source for stimulation. For repeat stimulation experiments, we used a rest period of ∼2 minutes in between sessions. All imaging sessions were carried out at room temperature in ACSF. When imaging in Ca^2+^-free media, we switched our imaging buffer to Ca^2+^-free ACSF. When imaging in nomifensine or TTX, we spike the imaging ACSF with a known volume and concentration of a nomifensine or TTX solution to attain the target concentration of nomifensine or TTX. We incubate the culture for 10 minutes at room temperature before carrying out post-drug activity imaging.

### Data Analysis

Each imaging stack was processed with a custom-built MATLAB code that converts the raw movie stack into a ΔF/F_0_ stack and generates correlated pixel components that we identify as hotspots of activity. To compute ΔF/F_0_, we first convolve the fluorescence time series movie (that is, raw F values) with a 2D gaussian (σ = 0.5 pixels). To obtain F_0_, a leaky cumulative minimum is calculated for each pixel in F, followed by repeated lowpass filtering to converge on a smooth F_0_ that obeys the minima. ΔF is then calculated as F -F_0_. Next, we partition the image into correlated components by applying nonnegative matrix factorization (NNMF) to ΔF with sparsity and contiguity constraints. This partitions the image into largescale correlated regions (that is, into pixel clusters with correlated activity). See Figure S4 and Figure S10 for example NNMF components. For each NNMF component, ΔF/F_0_ is then calculated using the component’s clustered pixels. Heatmaps and timeseries of ΔF/F_0_ traces are then generated for each movie file using native MATLAB capabilities. For simplicity, we refer to ΔF/F_0_ as ΔF/F in most of our figures and text. All calculations were done in MATLAB 2020 using the code located at the following repository: https://github.com/davidackerman/nnmf. To compute centroids of hotspots, and to generate overlay images, we used Fiji._70_ Statistics: All statistical tests of significance (*p* values) were computed and reported from unpaired two-tailed *t* test. We used MATLAB’s built-in boxplot algorithm to display distributions for some of our data. The definitions for relevant features of the box plot are as follows: red line = median, edges of box: 25^th^ and 75^th^ percentile, top and bottom hash lines: minimum and maximum values of non-outlier data, red points: outlier data.

### Post-hoc immunocytochemistry and Airyscan super resolution imaging

After activity imaging, cultures were fixed using 4% paraformaldehyde in 1X PBS for 15 mins at room temperature and washed twice with 1X PBS. Next, cells were permeabilized/blocked in 1X PBS with 0.1% Triton X-100 (PBST) for 1 hour at room temperature. All primary antibodies were incubated overnight at 4°C and secondary antibodies for 1 hour at room temperature in 1X PBS containing 1% BSA. Primary antibodies used in this study include chicken anti-tyrosine hydroxylase (1:1000; TYH 6767979, Aves labs), mouse anti-tyrosine hydroxylase (1:2000; T2928, Millipore Sigma), rabbit anti-tyrosine hydroxylase (1:2000; ab112, Abcam), mouse anti-bassoon (1:2000; ab82958, Abcam), chicken anti-MAP2 (1:1000, NB300-213, Novus Biologicals), guinea pig anti-synaptobrevin-2 (1:1000; SySy 104 318, Synaptic Systems). TH-GFP signal (after activity imaging) was amplified using a rabbit anti-GFP (1:2000; Chromotek PABG1) antibody. Alexa fluor 488 and Alexa fluor plus 647-conjugated secondary antibodies to mouse and guinea pig (Invitrogen) were applied at 1:2000 dilutions and Brilliant Violet 421-conjugated secondary antibody to chicken (Jackson Immunoresearch laboratories) was added at 1:1000 dilutions. Super resolution microscopy was performed at the Janelia Imaging Facility on a Zeiss LSM 880 inverted confocal microscope equipped with a Plan-Apochromat 63X/ 1.4 NA oil objective in Airyscan mode using a 32-channel gallium arsenide phosphide photomultiplier tube (120 nm lateral resolutions). Dopaminergic neurons of interest from the activity images were located by using MatTek dish grid location in the reflection mode.

