## Supplementary-Figures for "Visualizing Synaptic Dopamine Efflux with a 2D Nanofilm"

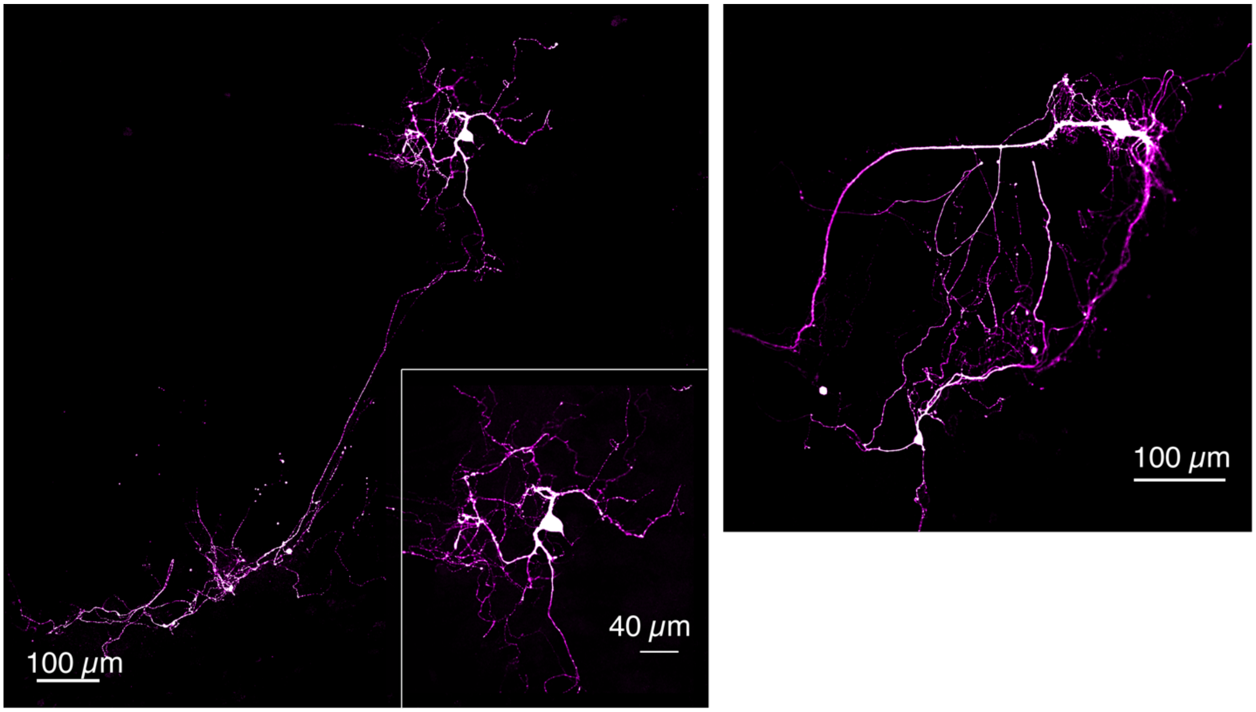


**Figure S1:** TH-immunofluorescence of Dopamine neurons grown on DopaFilm. Examples of TH+ dopamine neurons grown in co-culture with cortical neurons.

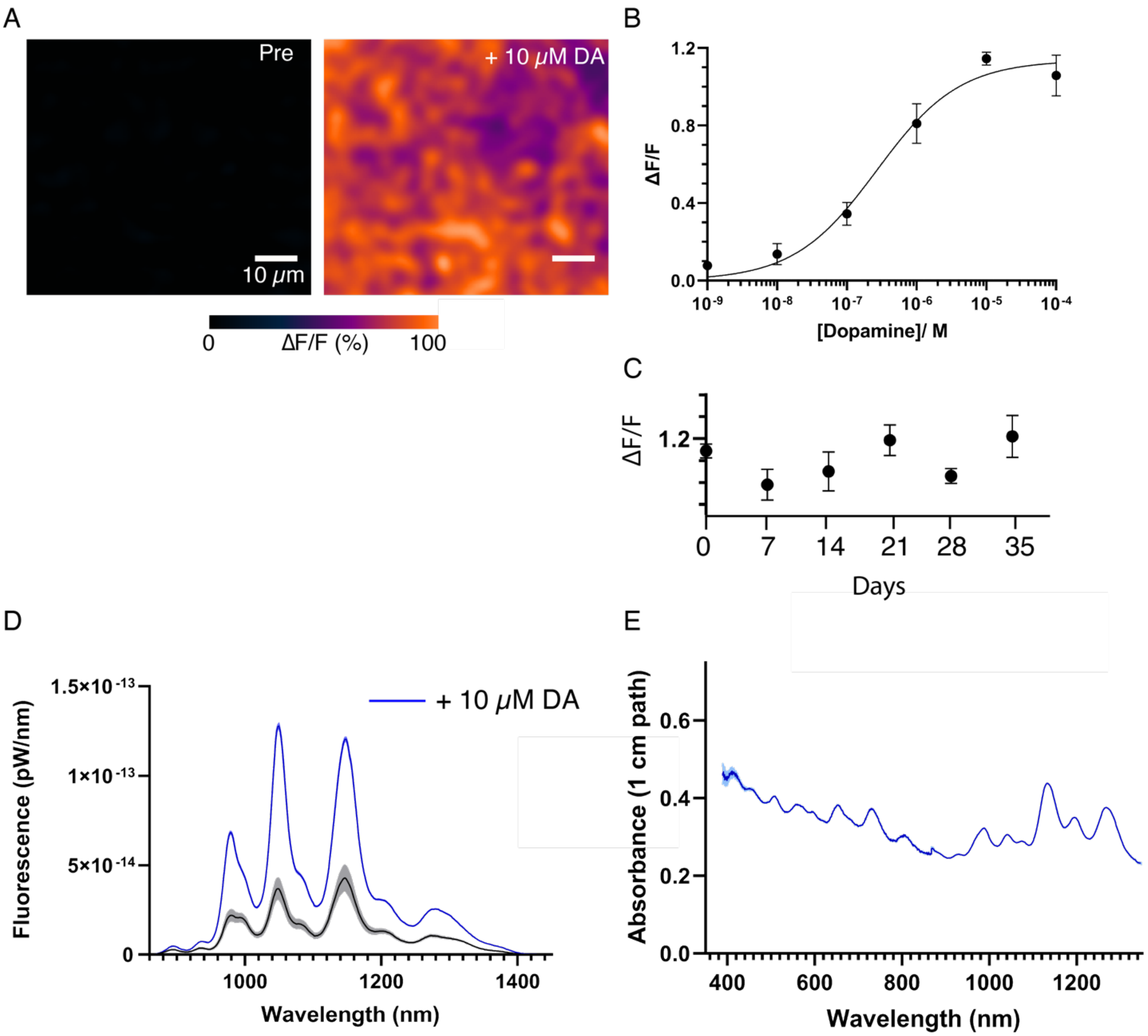


**Figure S2:** DopaFilm and solution phase nanosensor characterization.

(A) 10 μM of exogenous dopamine (DA) wash on DopaFilm shows diffuse ΔF/F response from DopaFilm substrate.
(B) Dose-response curve of DopaFilm. Each experimental data point has an error bar depicting standard deviation from different DopaFilm preparations (n = 3). Experimental data points were fit to the Hill equation (least-squares regression, black trace) to estimate an apparent dissociation constant ($K_d = 268\ nM$) and Hill coefficient ($n = 0.73$).
(C) DopaFilm response to 10 μM DA remains stable over 35 days. Each experimental data point has an error bar depicting standard deviation from different DopaFilm preparations (n = 3). x-axis: days post initial preparation.
(D) Solution phase fluorescence emission spectra of dopamine nanosensors, prepared from a conjugation of a multi-chiral single wall carbon nanotubes functionalized with single strand oligonucleotides, before (black trace) and after (blue trace) application of 10 μM DA. Bands indicate standard deviation from n = 3 measurements.
(E) Solution phase absorption spectrum of nanosensors.

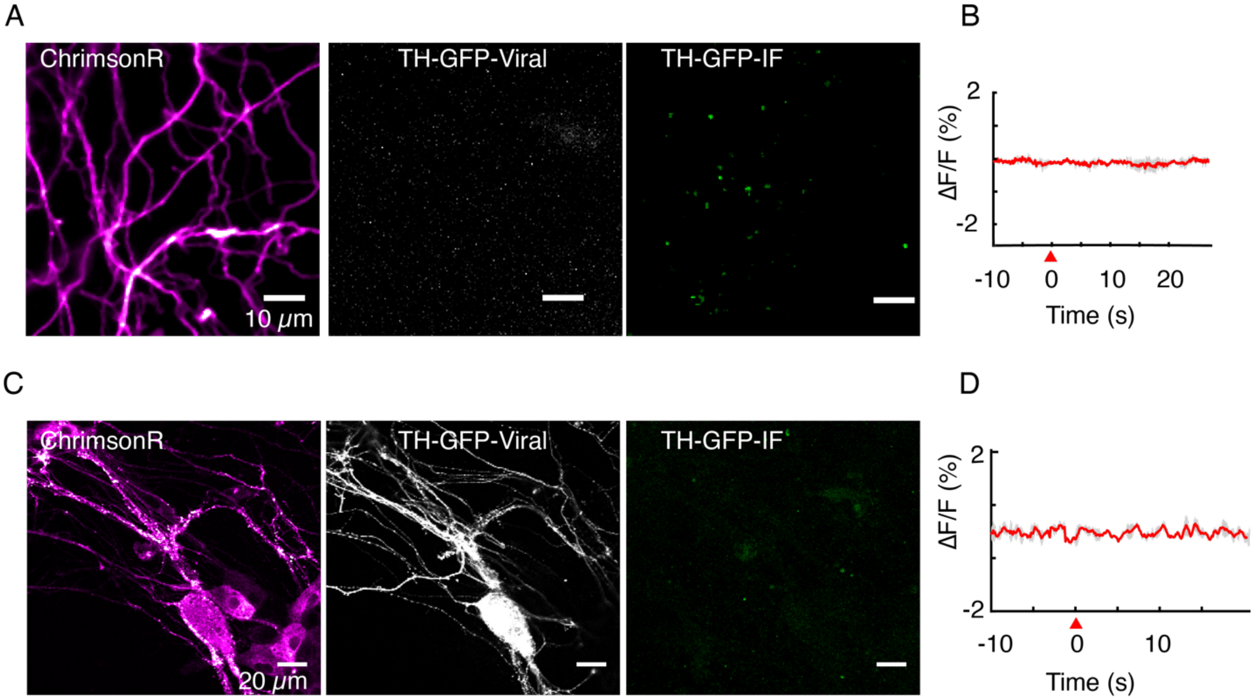


**Figure S3:** TH immunofluorescence is used to verify identity of putative dopamine neurons.

(A, B) Optical stimulation of ChrimsonR-positive and TH-GFP-negative axonal arbor elicits no evoked activity (n = 25 cells). Immunofluorescence (TH-GFP-IF) indicated no dopaminergic processes in the field of view.
(C, D) Optical stimulation of ChrimsonR-positive and TH-GFP-positive soma and dendritic arbor elicits no evoked activity (n = 10 cells). Retrospective immunofluorescence (TH-GFP-IF) indicates there are no dopaminergic processes in the field of view.
Red wedge: time of application of optical stimulation. Data in (B) and (D) are averaged from triplicate stimulations for FOVs shown in (A) and (C) respectively, and are not averages from the entire data set.

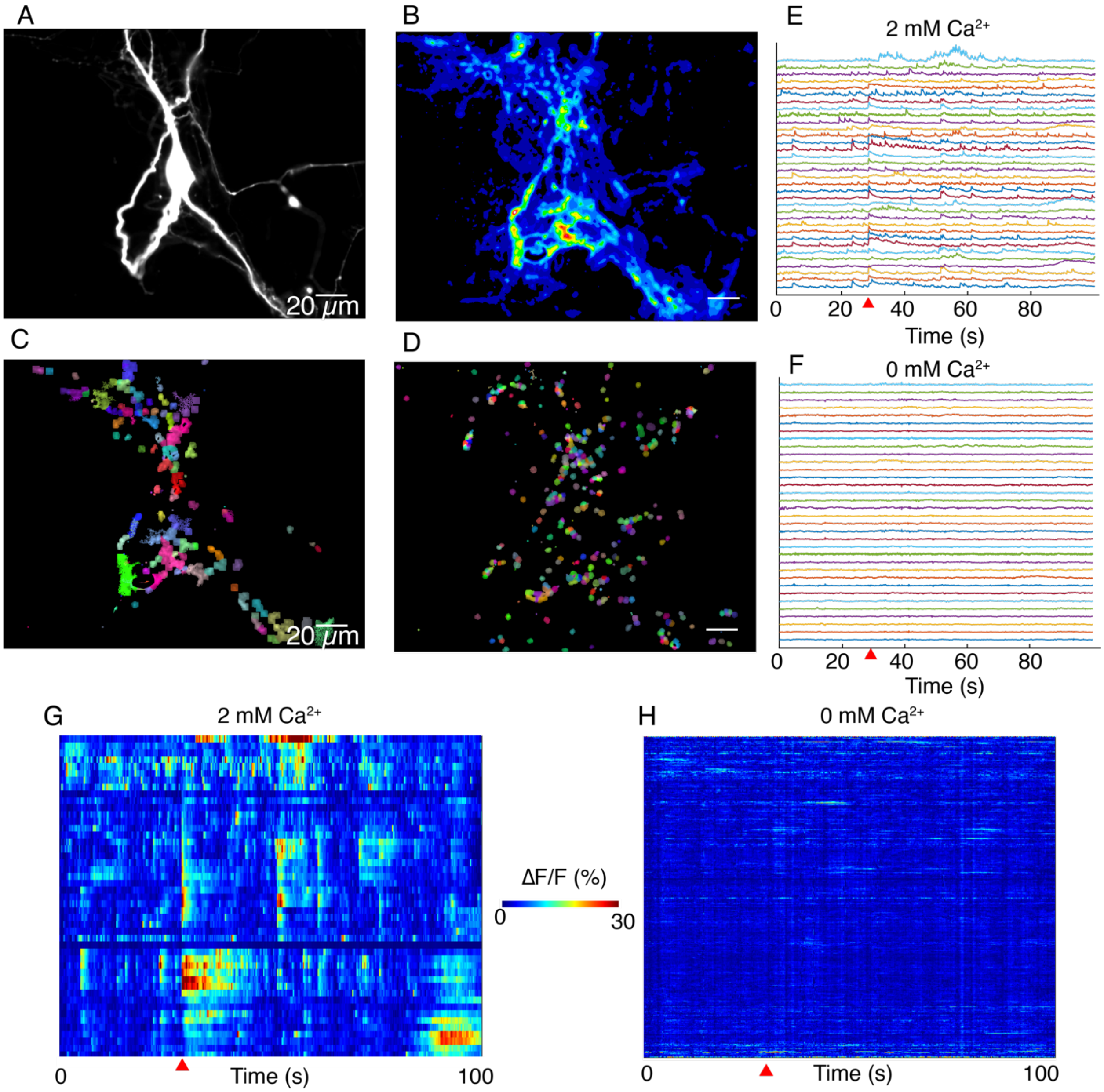


**Figure S4:** DopaFilm activity imaging in normal imaging buffer (ACSF with 2 mM $Ca^{2+}$) and buffer with no extracellular $Ca^{2+}$ ($Ca^{2+}$-free ACSF).

(A) TH-GFP image of a dopamine neuron.
(B) Maximum intensity projection from a 1000-frame ΔF/F movie of evoked and spontaneous activity imaging of neuron shown in (A), in normal ACSF.
(C) Non-negative matrix factorization (NNMF) decomposition applied to the ΔF/F movie in normal ACSF shows local clusters of pixels with highly correlated activity, color coded for better visualization. Similar colors in a local cluster of pixels corresponds to correlated activity.
(D) NNMF decomposition of movie stack of imaging in $Ca^{2+}$-free ACSF.
(E, F) ΔF/F time traces for a subset of the NNMF clusters (i.e., decomposed elements) from (C) and (D) respectively, off-set in y-axis for better visualization.
(G, H) Heat-maps of activity for all the NNMF components in (C) and (D) respectively.

Red wedge in figures (E)-(H) = time of optical stimulus application.

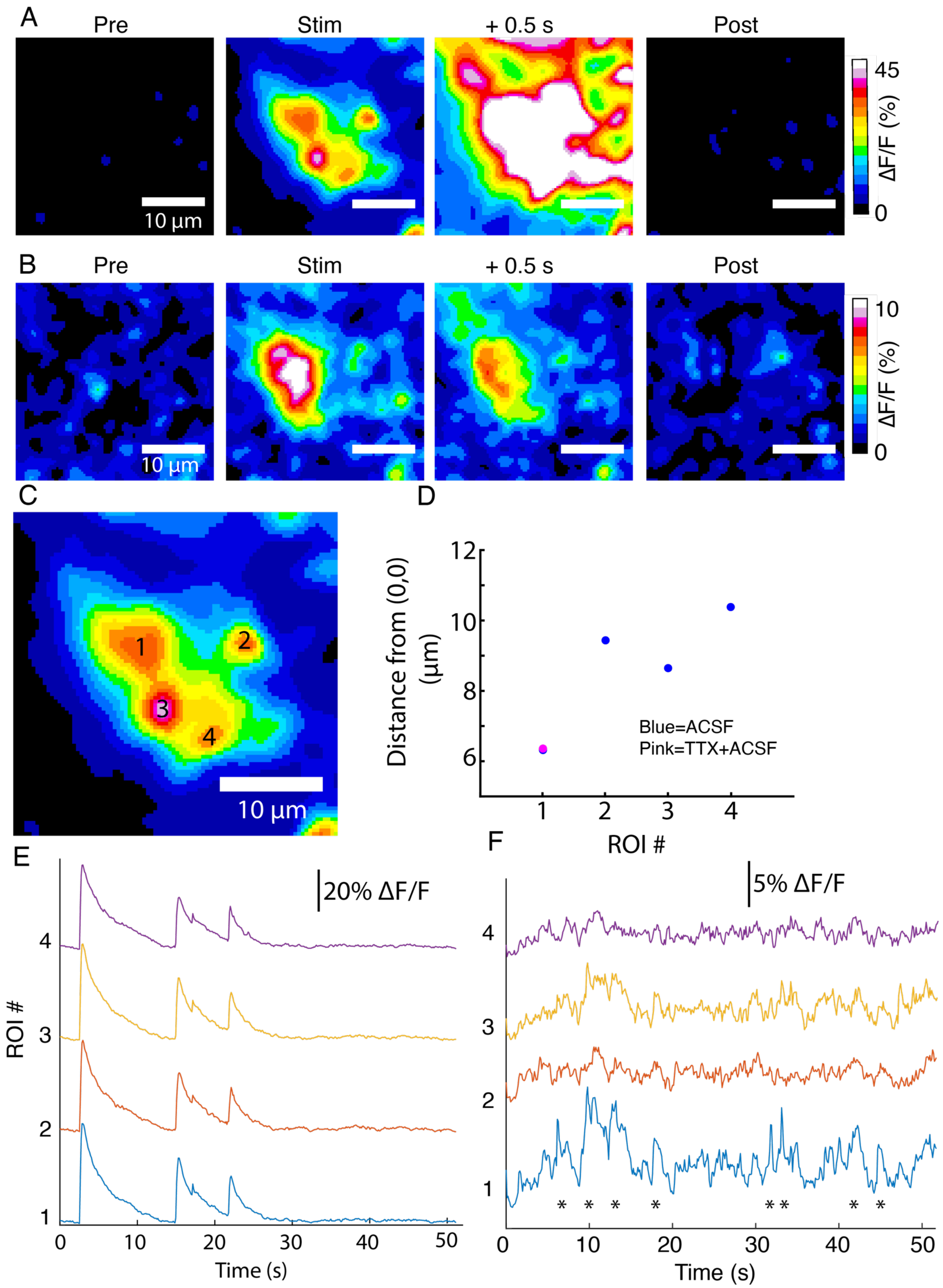


**Figure S5:** DopaFilm activity imaging in an axonal arbor in:

(A) normal ACSF
(B) ACSF with 10 $\mu$M of TTX.
(C) DopaFilm activity is driven by four putative release sites when imaging in ACSF, but activity is only seen in ROI #1 when TTX is applied.
(D) Centroid of hotspots in ACSF and ACSF+TTX. For ROI 1, centroids nearly overlap. For ROI2, 3 and 4, TTX centroids are missing.
(E, F) ΔF/F activity traces for ROIs 1 to 4 in ACSF (E) and ACSF+TTX (F).

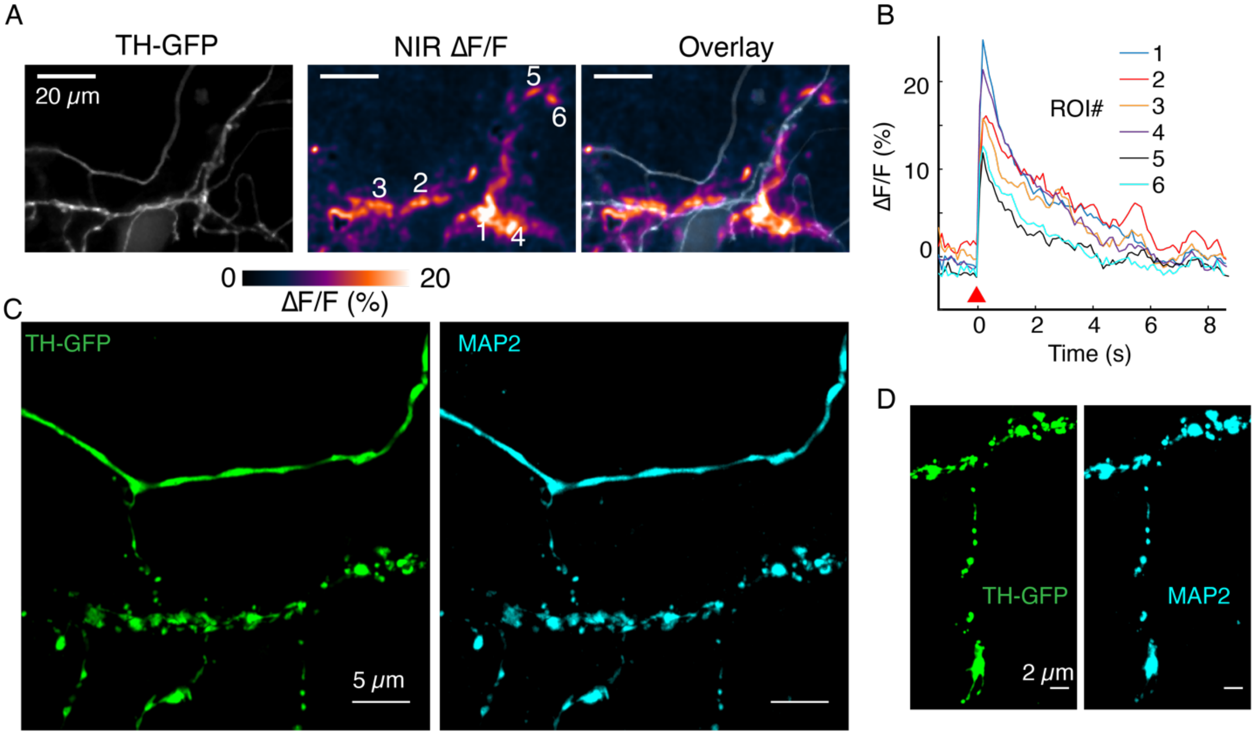


**Figure S6:** MAP2+ immunofluorescence demonstrates dendritic nature of processes.

(A) TH-GFP, NIR ΔF/F and overlay image corresponding to data in Figure 5D.
(B) Activity traces corresponding to ROIs 1 to 6 noted in (A).
(C) TH-GFP and MAP2 double- stain and super resolution image of FOV in (A).
(D) TH-GFP and MAP2 double stain of FOV in Figure 5B.

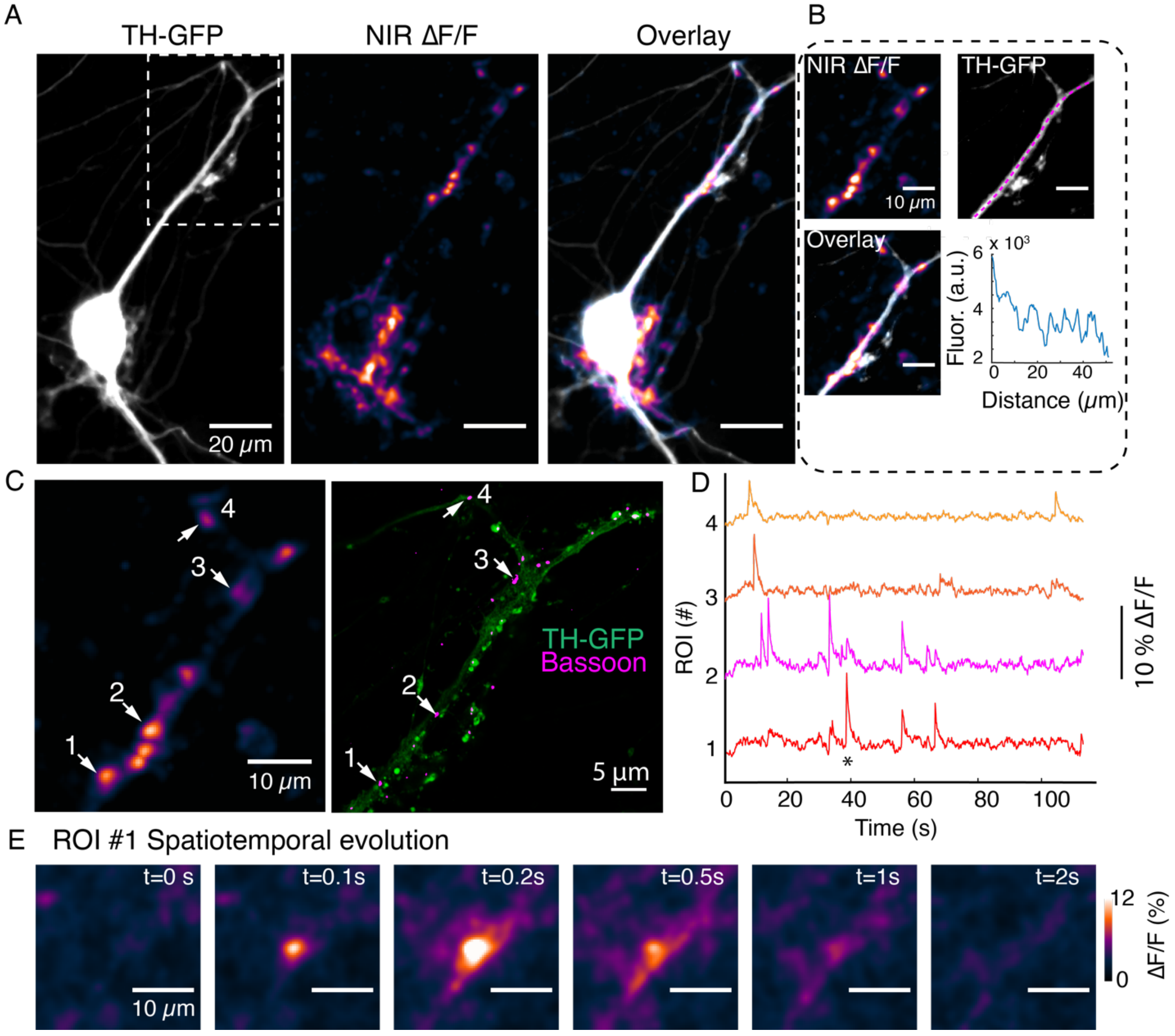


**Figure S7:** DopaFilm activity imaging at a dendrite of autonomously spiking dopamine neuron.

(A) TH-GFP image of a dopamine neuron, and maximum intensity projection from ΔF/F stack and overlay.
(B) Segment of activity in (A), (white box) and its TH-GFP intensity profile along the dendritic process (along magenta line), show no putative boutons.
(C) DopaFilm hotspot activity for the selection depicted in (A) and corresponding TH-GFP and Bassoon super resolution images. Bassoon puncta can be assigned to DopaFilm hotspot activity.
(D) ΔF/F traces for ROIs 1 to 4 shown in (C).
(E) Spatial dynamics of ROI#1 at the time depicted with * in the ROI1 trace shown (D)

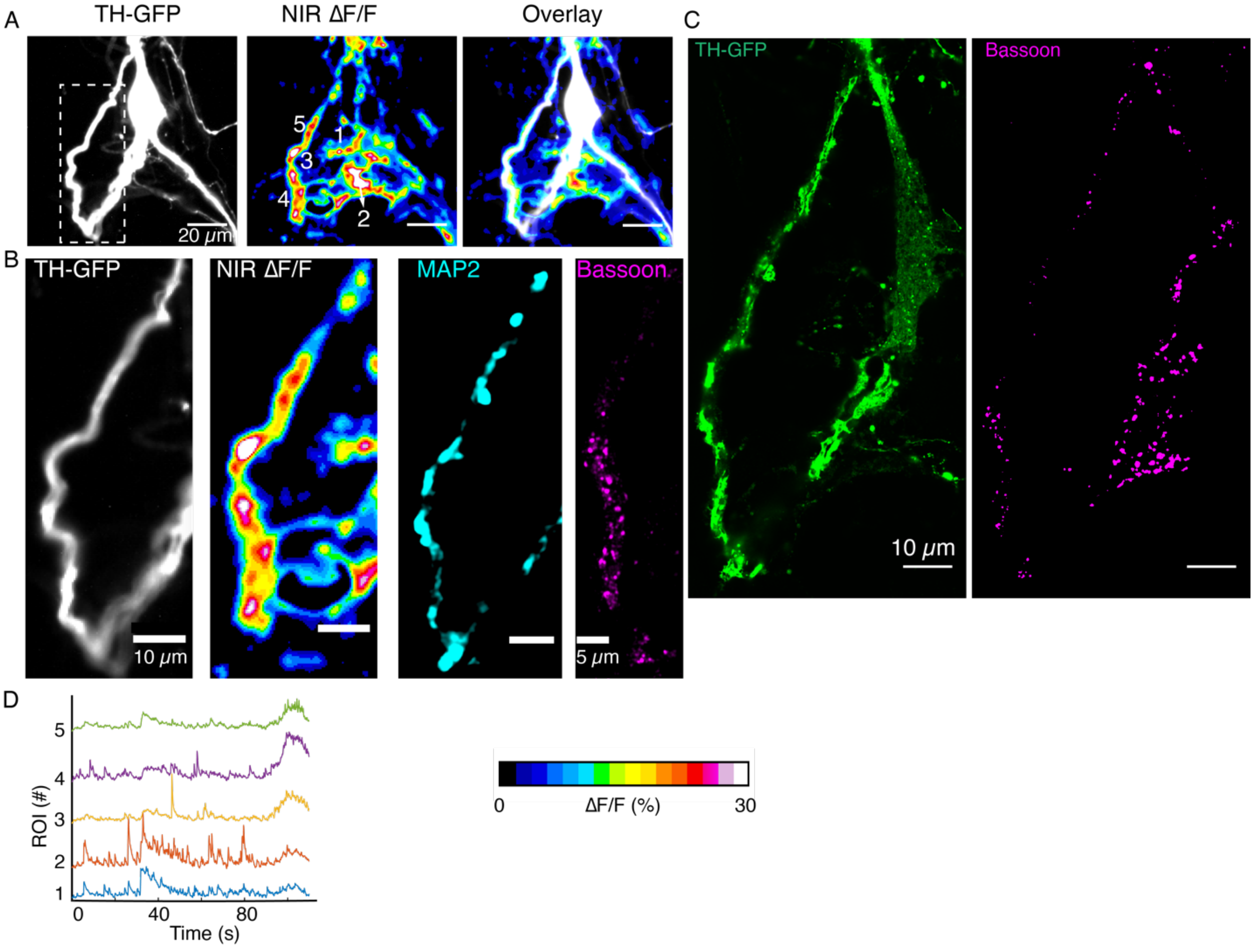


**Figure S8:** DopaFilm activity imaging of a dopamine neuron.

(A) TH-GFP image of a dopamine neuron, and maximum intensity projection from ΔF/F stack and overlay
(B) TH-GFP for the white box depicted in (A) and its DopaFilm activity along the dendritic process, as well as MAP2 and Bassoon super-resolution images.
(C) TH-GFP and Bassoon super-resolution images for the bigger field of view in (A).
(D) ΔF/F time traces for ROIs 1 to 5 depicted in (A) NIR ΔF/F panel.

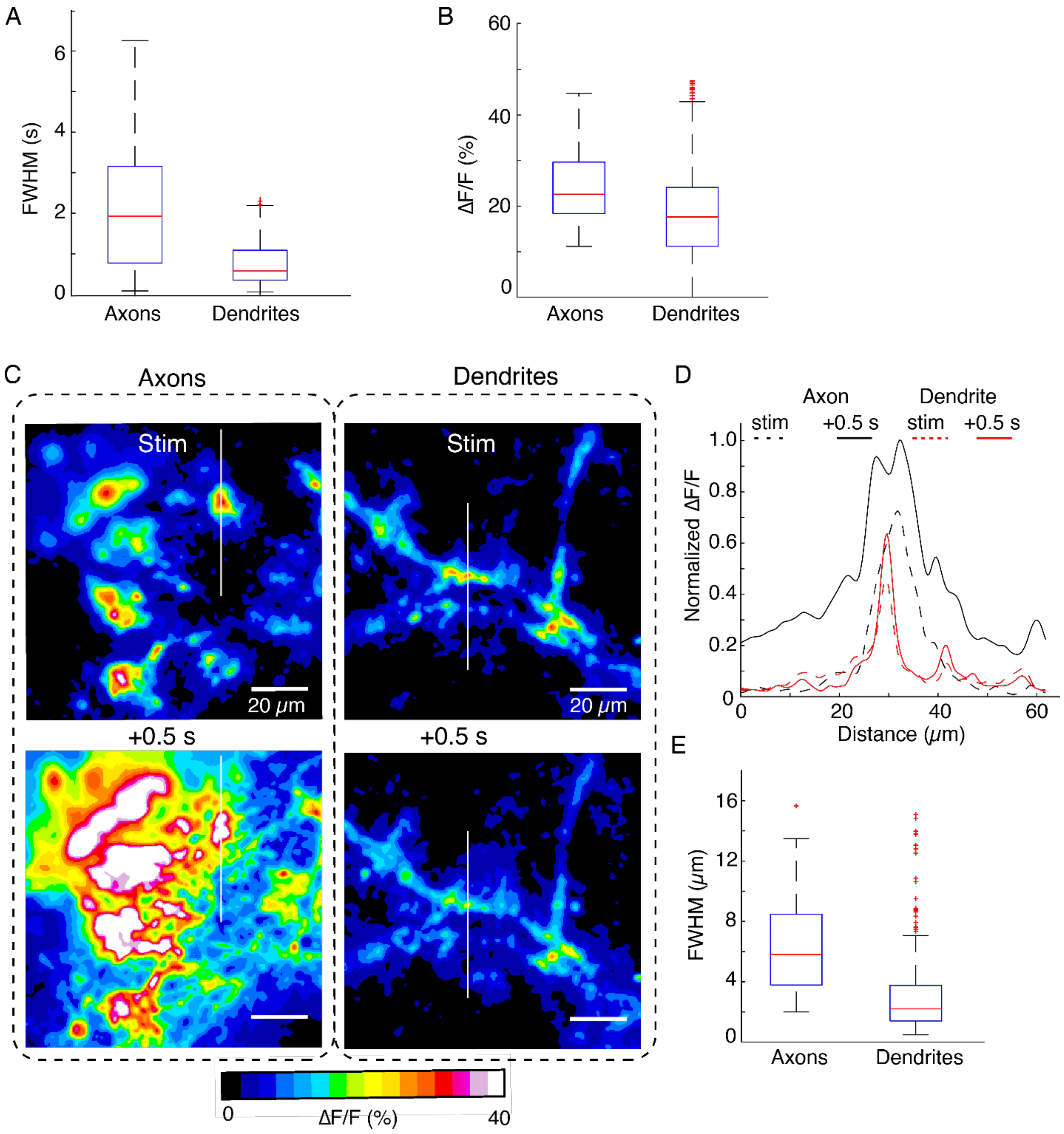


**Figure S9** Comparison of spatiotemporal dynamics of DopaFilm hotspots in axonal arbors and dendritic processes after optical stimulation (evoked).

(A, B) Box plots comparing pooled data for temporal full width at half-max (FWHM) (s) of transient traces and peak ΔF/F in axons and dendrites. Here, FWHM is defined in the time domain as a measure of how long (in seconds) the transients persist before diffusive and dopamine transporter (DAT)-mediated clearance. Mean ± SD: Peak ΔF/F (%) in axons: 25 ± 8.3. In dendrites: 18 ± 9.3. Unpaired t-test: $p < 10^{-4}$. Temporal FWHM (s) in axons: 1.75 ± 1.2. In dendrites: 0.9 ± 0.7. Unpaired t-test: $p < 10^{-4}$.
(C) Spatial spread of DopaFilm activity in an axonal arbor (left column) and dendritic arbor (right column). ΔF/F images are shown immediately following stimulation (stim, top row) and 0.5 seconds after stimulation

(+0.5 s, bottom row). Notice the diffusive broadening of the signal in the axons, which occurs only to a limited extent in dendrites.

(D) Line profiles along the white lines depicted on the images in (C). Dash lines represent profile immediately following stimulation (stim) and solid lines represent line profiles after 0.5 s has elapsed. ΔF/F is normalized to the peak ΔF/F observed in the axonal arbor. Notice how activity in axons has a bigger spatial extent than those in dendrites.

(E) Spatial FWHM ($\mu$m) in axons vs. dendrites. Mean $\pm$ SD; axons: 6.6 $\mu$m $\pm$ 3.6 $\mu$m; dendrites: 3.2 $\mu$m $\pm$ 3$\mu$m. Unpaired t-test: $p < 10^{-4}$. Data is from n = 32 dendritic processes pooled from n = 12 dopamine neurons. For axons, we pooled data from n = 6 axonal arbors.

Boxplot definitions: red line = median, edges of box: 25th and 75th percentile, top and bottom hash lines: minimum and maximum values of non-outlier data, red points: outlier data.

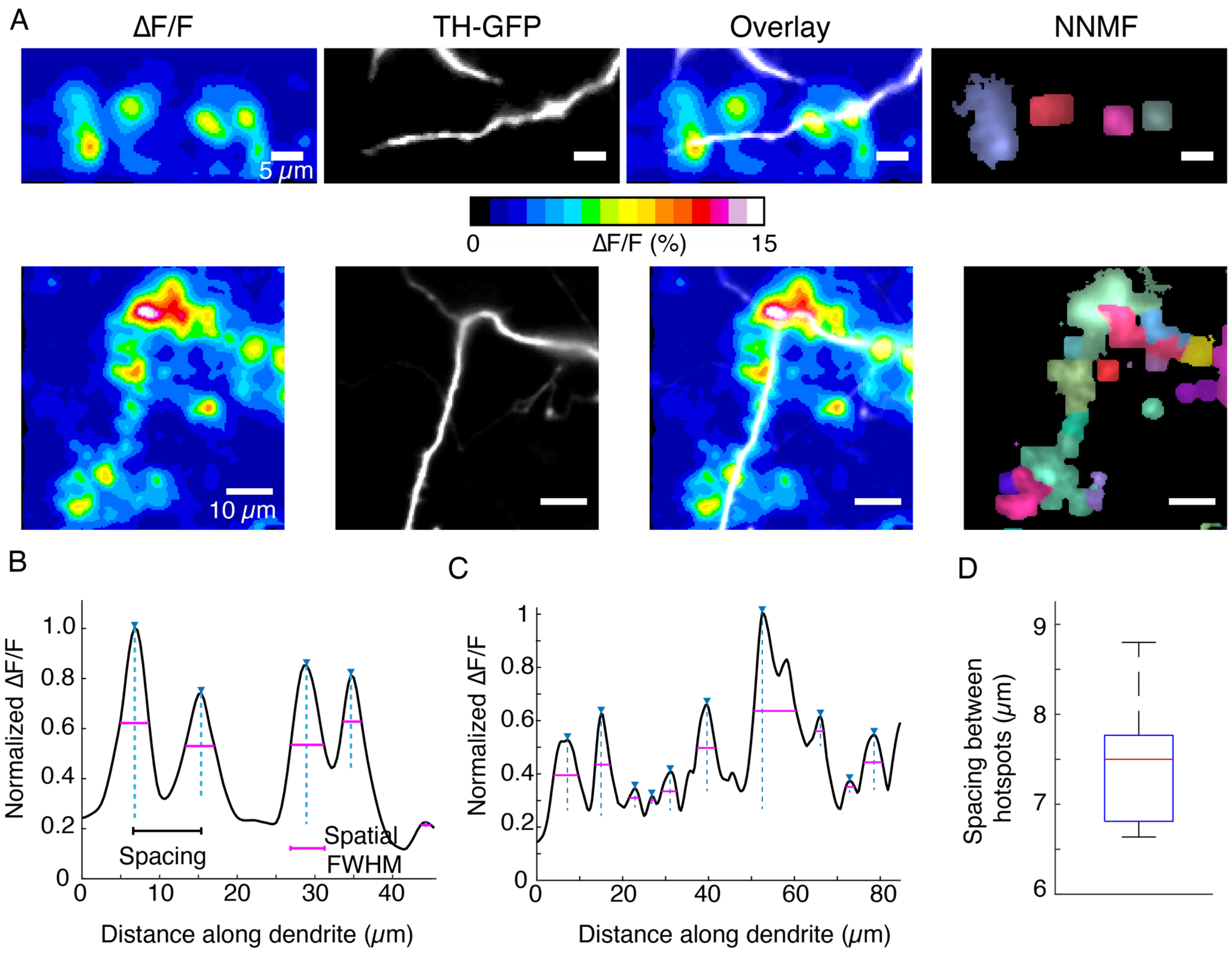


**Figure S10:** DopaFilm hotspot activity along dendritic processes.

(A) Top and bottom rows: ΔF/F, TH-GFP and overlay. Right-most panels: NNMF decomposition of ΔF/F stack identifies clusters of pixels with highly correlated activity, consistent with the ΔF/F hotspots.

(B, C) Line profiles of ΔF/F computed along the profile of the dendrites corresponding to the top and bottom rows in (A), respectively. Custom code identifies peaks, distance between peaks (spacing, black line segment) and the spatial extent of each signal, approximated by the FWHM (magenta line segment).

(D) Box plot of spacing between hotspots observed on dendritic process. Box plot distribution for FWHM ($\mu$m) is shown in Figure S9.

Data is pooled from n = 32 dendritic processes pooled from n = 12 dopamine neurons. Boxplot definitions: red line = median, edges of box: 25th and 75th percentile, top and bottom hash lines: minimum and maximum values of non-outlier data.

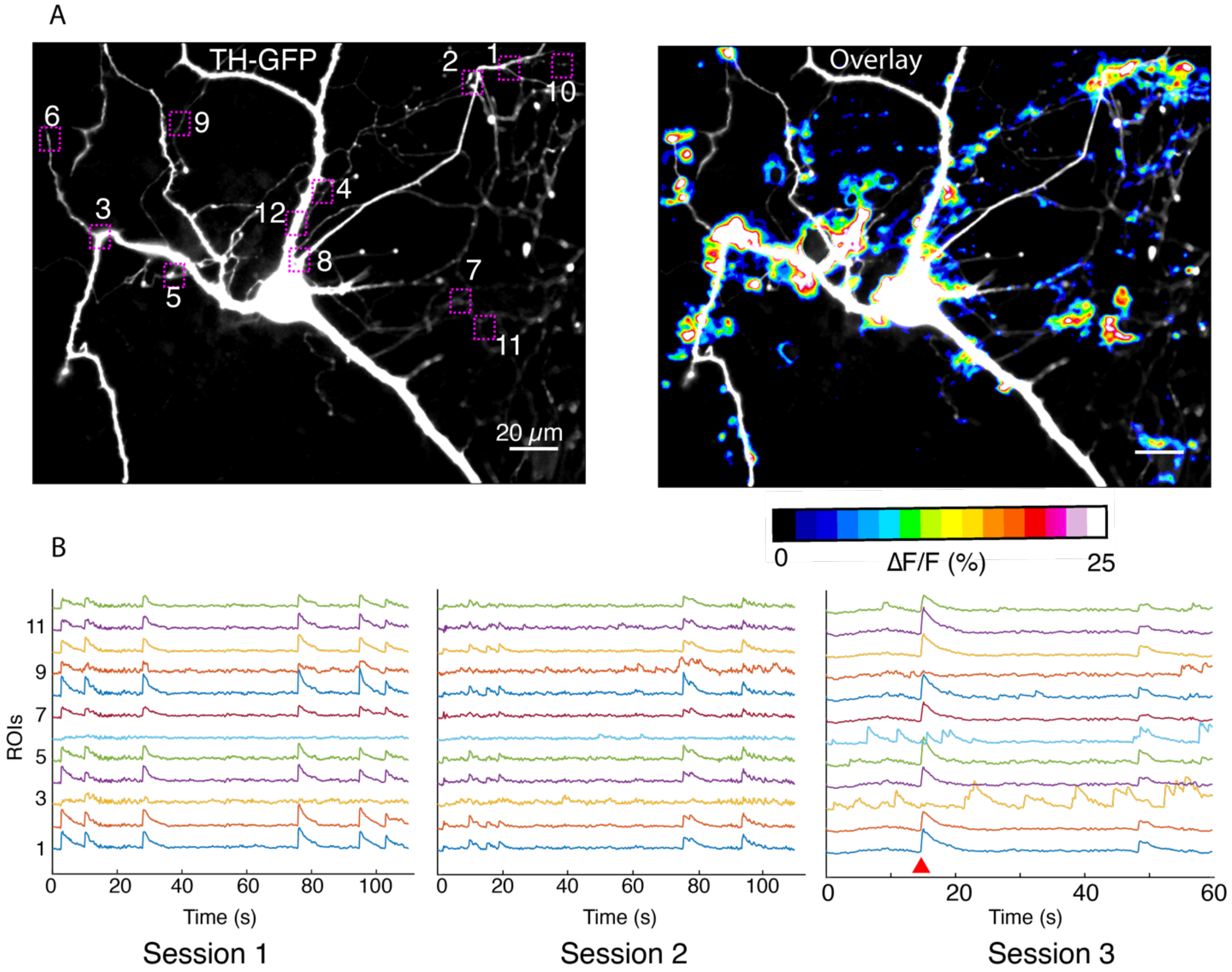


**Figure S11:** Imaging in dendritic processes of a dopamine neuron.

(A) TH-GFP image and its overlay with evoked ΔF/F DopaFilm activity image.
(B) ΔF/F traces for the ROI boxes numbered in (A) from n = 5 imaging session (n = 3 sessions shown here). Note that most activity is driven by autonomous spiking except where the red wedge indicates optical stimulus driven activity (in session #3).

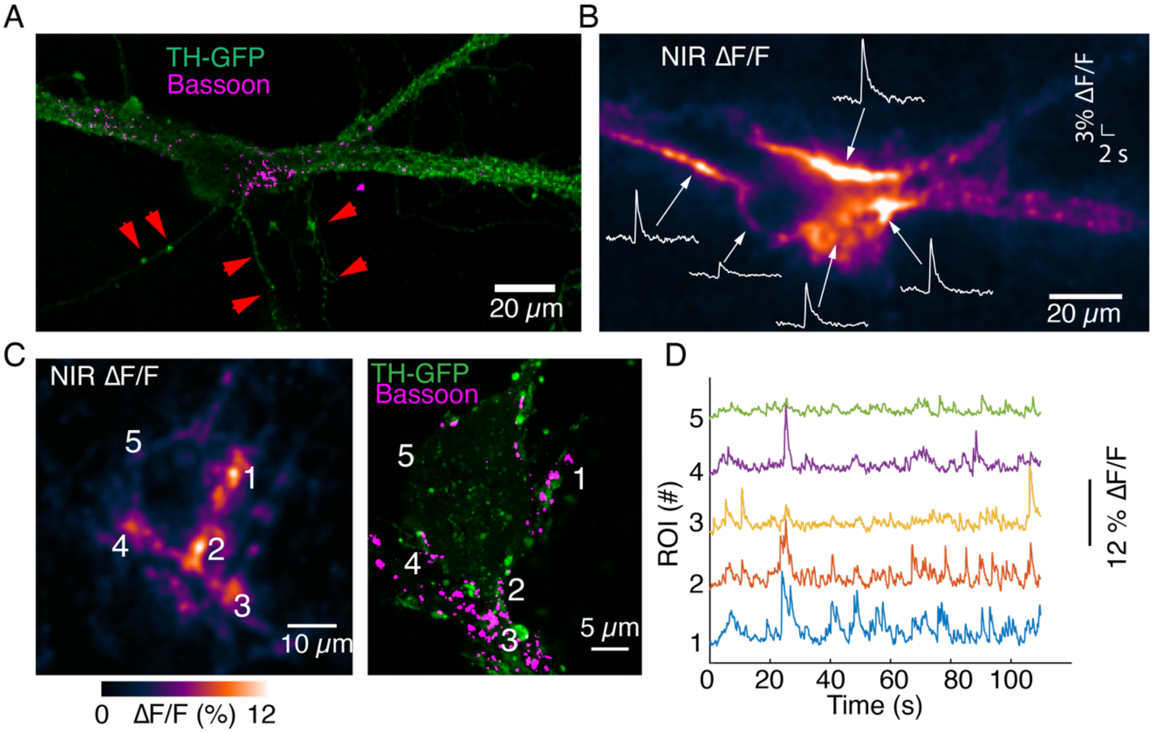


**Figure S12:** Imaging activity from proximal dendrites associated with the soma of dopamine neurons.

(A) TH-GFP image and Bassoon puncta on the major dendrites of a dopamine neuron.
(B) ΔF/F activity evoked by optical stimulation. Notice activity arising from major dendritic trunks and at the junction of soma and major dendrites. Also notice lack of activity in processes that lack Bassoon puncta (red arrows).
(C) Autonomous spiking activity around the soma of a dopamine neuron. Max ΔF/F projection is shown on left. Notice activity at the major dendrite leaving the cell body (around area depicted as ROI #3) and comingling dendrites (around area depicted as ROI #1, ROI#4), both enriched in Bassoon puncta. ROI#5 exhibits reduced activity relative to ROIs #1 to #4 and has lower levels of Bassoon expression.
(D) Activity traces corresponding to ROIs depicted in (C).

**Movie S1:** Imaging in axons.
**Movie S2:** Imaging in axons before application of TTX.
**Movie S3:** Imaging in axons after application of TTX.
**Movie S4:** Evoked release from a dendritic process.
**Movie S5:** Spontaneous activity from a dendritic process.
**Movie S6:** Dendritic activity around soma of dopamine neuron.
