## Supplementary figures and images for "Visualizing Synaptic Dopamine Efflux with a 2D Nanofilm"

### Movie-S1

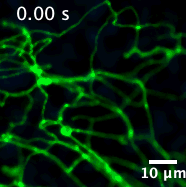

### Movie-S2

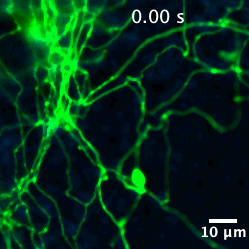

### Movie-S3

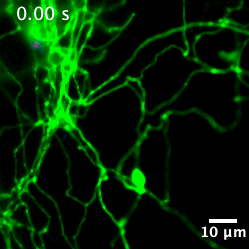

### Movie-S4

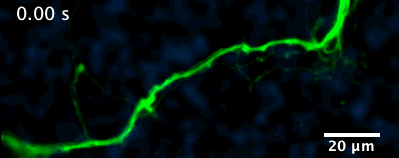

### Movie-S5

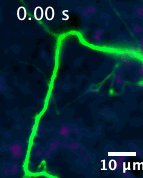

### Movie-S6

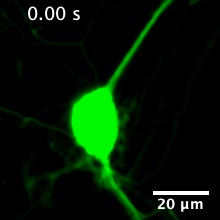
